# Leveraging gene co-regulation to identify gene sets enriched for disease heritability

**DOI:** 10.1101/2021.07.22.453442

**Authors:** Katherine M. Siewert-Rocks, Samuel S. Kim, Douglas W. Yao, Huwenbo Shi, Alkes L. Price

**Affiliations:** Department of Epidemiology, Harvard T.H. Chan School of Public Health, Boston, MA, USA; Program in Medical and Population Genetics, Broad Institute of MIT and Harvard, Cambridge, MA, USA; Department of Electrical Engineering and Computer Science, Massachusetts Institute of Technology, Cambridge, MA, USA; Program in Systems, Synthetic, and Quantitative Biology, Harvard University, Cambridge, MA, USA; Department of Biostatistics, Harvard T.H. Chan School of Public Health, Boston, MA, USA

## Abstract

Identifying gene sets that are associated to disease can provide valuable biological knowledge, but a fundamental challenge of gene set analyses of GWAS data is linking disease-associated SNPs to genes. Transcriptome-wide association studies (TWAS) can be used to detect associations between the genetically predicted expression of a gene and disease risk, thus implicating candidate disease genes. However, causal disease genes at TWAS-associated loci generally remain unknown due to gene co-regulation, which leads to correlations across genes in predicted expression. We developed a new method, gene co-regulation score (GCSC) regression, to identify gene sets that are enriched for disease heritability explained by the predicted expression of causal disease genes in the gene set. GCSC regresses TWAS chi-square statistics on gene co-regulation scores reflecting correlations in predicted gene expression; GCSC determines that a gene set is enriched for disease heritability if genes with high co-regulation to the gene set have higher TWAS chi-square statistics than genes with low co-regulation to the gene set, beyond what is expected based on co-regulation to all genes. We verified via simulations that GCSC is well-calibrated, and well-powered to identify gene sets that are enriched for disease heritability explained by predicted expression. We applied GCSC to gene expression data from GTEx (48 tissues) and GWAS summary statistics for 43 independent diseases and complex traits (average *N* =344K), analyzing a broad set of biological pathways and specifically expressed gene sets. We identified many enriched gene sets, recapitulating known biology. For Alzheimer’s disease, we detected evidence of an immune basis, and specifically a role for antigen presentation, in analyses of both biological pathways and specifically expressed gene sets. Our results highlight the advantages of leveraging gene co-regulation within the TWAS framework to identify gene sets associated to disease.

## Introduction

Gene set enrichment analyses of GWAS data can provide valuable insights about disease etiology [1]. MAGMA [2] and other widely used methods [3–5] assess enrichment of disease signal for SNPs located near genes in a gene set, but linking SNPs to genes via proximity is imprecise; in particular, recent studies have estimated that the gene closest to a lead GWAS SNP is the causal disease gene less than half of the time [6–9]. Gene expression data provides a more precise way to link SNPs that are eQTLs to genes whose expression they regulate. For example, MESC [10] estimates the disease heritability mediated by predicted gene expression, and can also be used to identify gene sets enriched for disease heritability mediated by predicted gene expression. However, MESC relies on strong modelling assumptions and does not pool information across different eQTLs for the same gene to leverage directional consistency (i.e. consistency in the product of eQTL and GWAS effect directions between SNPs).

Here, we introduce gene co-regulation score (GCSC) regression, a new method that identifies gene sets enriched for disease heritability explained by predicted expression. GCSC makes use of the transcriptome-wide association study (TWAS) framework [6, 9, 11], which uses eQTL effects from gene expression prediction models and summary statistics from GWAS to test genes for association between predicted gene expression and disease; this enables GCSC to pool information across different eQTLs for the same gene to leverage directional consistency. TWAS-associated genes may not be biologically causal due to correlation in predicted expression between genes (gene co-regulation), which can arise due to shared causal eQTLs or linkage disequilibrium (LD) between eQTLs [9, 12]. GCSC leverages gene co-regulation to identify enriched gene sets (analogous to stratified LD score regression [13], which leverages LD between SNPs to identify enriched SNP sets).

We validate the calibration and power of GCSC via extensive simulations and find it outperforms MESC. We then apply GCSC to GWAS summary statistics for 43 independent diseases and complex traits (average *N* =344K) in conjunction with gene expression data from 48 GTEx (v7) tissues [7], analyzing 38 non-disease-specific gene sets, 11,394 biological pathways, and 262 specifically expressed gene sets. We discover both established and more novel gene set enrichments for a variety of traits, demonstrating the power of GCSC to uncover important gene sets for disease.

## Results

### Overview of methods

GCSC relies on the TWAS chi-square association statistics of each gene with disease [6, 11], which include both the effect of the focal gene and the effects of co-regulated genes (defined here as genes with correlated values of predicted expression) [9]. GCSC assesses the contribution of a gene set to disease heritability explained by predicted expression by regressing TWAS chi-square statistics on two gene co-regulation scores, reflecting co-regulation with all genes and co-regulation with genes in the gene set, respectively. GCSC determines that a gene set is enriched for disease heritability if genes with high co-regulation to the gene set have higher TWAS chi-square statistics than genes with low co-regulation to the gene set, beyond what is expected based on co-regulation to all genes.

In detail, the expected TWAS *χ*^2^ statistic for gene *g* is

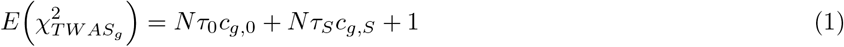

where *N* is GWAS sample size, *c*_*g*,0_ is the co-regulation score of gene *g* with respect to all genes (defined as 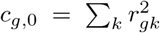, where 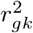 is the squared correlation in predicted expression of gene *g* and gene *k*), *c*_*g,S*_ is the co-regulation score of gene *g* with respect to genes in gene set *S* (defined as 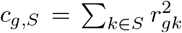), *τ*_0_ is the per-gene heritability explained by predicted expression of all genes without including the excess heritability contributed by genes in gene set *S*, and *τ*_*S*_ is the per-gene excess heritability contributed by genes in gene set *S*. We estimate the heritability explained by predicted gene expression without including the excess heritability contributed by genes in gene set *S* as 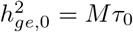, and estimate the excess heritability contributed by genes in gene set *S* as 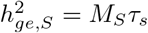, where *M* is the total number of genes and *M*_*S*_ is the number of genes in gene set *S*. Thus, the total heritability explained by predicted gene expression is 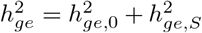. Further details (including the use of regression weights to increase power, and a correction for the bias arising from the distinction between predicted expression and the total cis-genetic component of expression) are provided in the Methods section, and a derivation of Equation 1 is provided in the Supplementary Note.

Eq. (1) allows us to estimate the excess heritability contributed by genes in gene set 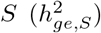 via a bilinear regression of TWAS chi-square statistics on *c*_*j*,0_ and *c*_*j,S*_ (Fig. 1), with free intercept (see Methods). A 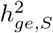 value that is significantly different from zero indicates a gene set that is enriched or depleted for heritability; we estimate standard errors via genomic block-jackknife, analogous to previous work [13]. GCSC can be applied to either binary gene sets or continuous-valued gene scores and can be used to jointly test for conditional heritability enrichment of multiple gene sets (or gene scores), analogous to the application of stratified LD score regression [13] (S-LDSC) to SNP annotations. GCSC can be applied either to TWAS chi-square statistics (and co-regulation scores) for a single tissue, or by concatenating TWAS chi-square statistics (and co-regulation scores) for multiple tissues (see Methods). We have publicly released open-source software implementing the GCSC method (see Code Availability).

**Figure 1:**
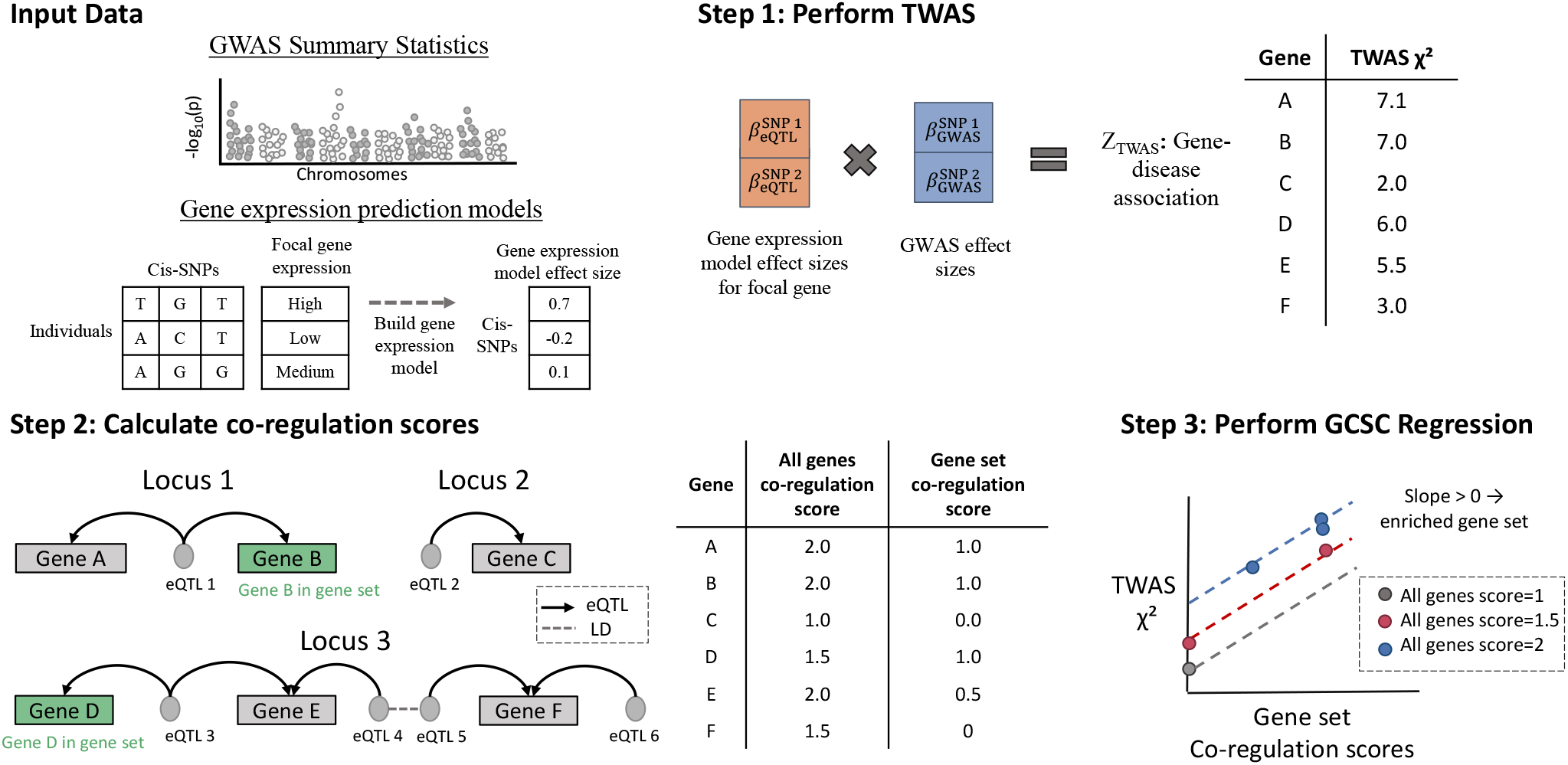
GCSC Schematic. (1) TWAS is performed using previously constructed gene expression prediction models. (2) Co-regulation scores (which reflect the amount of cis-genetically determined gene expression tagged by the prediction expression of each gene) are calculated using the gene expression prediction models; co-regulation can be due to shared eQTLs (solid lines) or LD between eQTLs (dashed line). (3) GCSC regression is performed to assess enrichment for disease heritability explained by predicted expression of genes in the gene set. In this example, within the same All genes co-regulation score bin, genes with a higher gene set co-regulation score have higher TWAS *χ*^2^ (i.e. the slope denoted by the dotted line is positive), indicating heritability enrichment.

We obtained gene expression prediction models (see URLs) that were constructed using the FUSION package [6] applied to GTEx gene expression data [7] (v7; 48 tissues, 18,094 protein-coding genes). We retained those with significant cis-heritability of gene expression (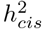, estimated using GCTA [14]) in each respective tissue (average sample size=214, average number of significantly cis-heritable genes=4,834) (Supplementary Table 1). We used the gene expression prediction models to compute TWAS *χ*^2^ statistics (using FUSION [6]) for 43 independent diseases and complex traits with publicly available GWAS summary statistics (average *N* =344K; Supplementary Table 2), and to compute co-regulation scores in each tissue (see Methods). We applied GCSC to evaluate the heritability enrichment of 38 non-disease-specific gene sets, 11,394 biological pathways and 262 sets of specifically expressed genes. We applied GCSC by concatenating data from all 48 tissues, unless otherwise noted. We have publicly released all TWAS chi-square statistics, co-regulation scores, and GCSC output produced in this study (see Data Availability).

### Evaluation of GCSC using simulations

We performed simulations using real genotypes and simulated gene expression and disease phenotypes to assess the robustness and power of GCSC to identify gene sets that are enriched for disease heritability explained by the predicted expression of causal disease genes in the gene set. We used individual-level data from European samples from the 1000 Genomes project [15] to simulate gene expression in two tissues (10,000 genes; 500 and 80 gene expression samples respectively). We used individual-level data from the UK Biobank [16])) to simulate disease phenotypes (see Methods). Default simulation parameters were set to a uniform distribution between 1 and 5 for the number of causal cis-eQTLs for each gene in each tissue, 12% for the cis-heritability of gene expression in each tissue, 75% for the cross-tissue cis-genetic correlation (consistent with estimates in [17]), and 10% for the disease heritability explained by the cis-genetic component of gene expression in each of the two tissues (summing to a total disease heritability of 20%, consistent with estimates of disease heritability mediated by gene expression in [10]); other parameter values were also explored. We compared GCSC to MESC [10], a method that estimates the enrichment of disease heritability mediated by gene expression of genes in a gene set. We did not evaluate S-LDSC [13] or MAGMA [2] in these simulations; those methods do not use eQTL data, and thus the relative performance of GCSC (and MESC) to those methods would be highly sensitive to assumptions about the role of gene expression in disease architectures. However, we include a comparison to those methods using real data in the next section.

We first evaluated the robustness of GCSC and MESC in null simulations with 1% of the 10,000 genes in the candidate gene set; null simulations refers to the scenario with nonzero disease heritability explained by gene expression, but no enrichment for genes in the candidate gene set (1x enrichment). We determined that both GCSC and MESC enrichment estimates were approximately unbiased (Fig. 2A, left, Supplementary Table 3). We also determined that both GCSC and MESC p-values were approximately well-calibrated (Fig. 2B, Supplementary Table 4).

**Figure 2:**
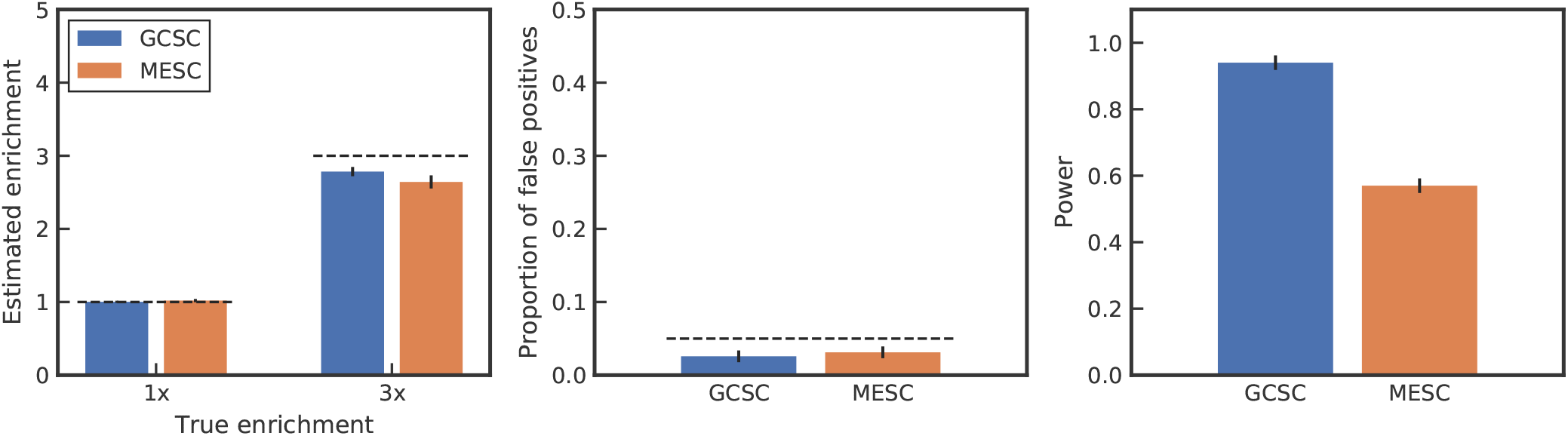
Performance of GCSC and MESC in simulations. We report (A) estimated enrichment in null simulations with no heritability enrichment in the gene set (1x) and causal simulations with heritability enrichment in the gene set (3x); (B) the proportion of null simulations in which each method produced a (1-sided) enrichment p-value less than 0.05; and (C) the proportion of causal simulations in which each method produced a (1-sided) enrichment p-value less than 0.05. Error bars denote 1 standard error. Numerical results are reported in Supplementary Table 3.

We next evaluated the robustness and power of GCSC and MESC in causal simulations with 1% of the 10,000 genes in the candidate gene set; causal simulations refers to the scenario with enrichment of disease heritability explained by gene expression for genes in the candidate gene set (3x enrichment). We determined that both GCSC and MESC enrichment estimates were slightly conservative (Fig. 2A, right). We also determined that GCSC was considerably more powerful than MESC (Fig. 2C). We attribute this to the greater flexibility of GCSC in (i) leveraging the TWAS framework, allowing GCSC to check for directional consistency between eQTLs and (ii) not assuming that per-gene cis-heritability of gene expression should be proportional to per-gene disease heritability explained by gene expression (an assumption that is made by MESC but was shown in [10] to be incorrect).

We performed 5 secondary analyses involving simulations at other parameter settings. First, we simulated the cis-heritability of gene expression and the disease heritability explained by gene expression to vary across genes (sampled from a normal distribution) instead of having a fixed value. This resulted in a reduction in power for both GCSC and MESC, but the power of GCSC remained significantly higher than MESC (Supplementary Fig. 1). Second, we simulated a scenario where in addition to the disease heritability of 20% mediated by gene expression, there is an additional disease heritability of 20% caused by SNP effects independent of eQTL effects. We determined that this scenario slightly decreased the power of both methods (Supplementary Fig. 2). Third, we simulated a wide variety of gene set sizes (0.5% to 40% of the 10,000 genes) and enrichment values, and determined that GCSC had approximately well-calibrated standard errors and higher power than MESC throughout this range (Supplementary Table 4, Supplementary Fig. 3 and 4). Fourth, we reduced the disease heritability explained by gene expression in each tissue from 10% to 5%. We determined that this reduced the power of both methods, as expected (Supplementary Fig. 5). Fifth, we used a GWAS sample size of 20,000 individuals instead of the default value of 50,000. We determined that this slightly decreased the power of both methods (Supplementary Fig. 6).

### Evaluation of GCSC in real data using random and positive-control gene sets

We further evaluated GCSC by applying it to GTEx gene expression data [7] (v7; 48 tissues, 18,094 protein-coding genes) and GWAS summary statistics for 43 disease and complex traits (average *N* =344K; Supplementary Table 2), analyzing both random gene sets and two well-studied “positive-control” gene sets, high-pLI genes [18] (2,791 genes) and Mouse Genome Informatics (MGI) essential genes [19] (2,240 genes) (Supplementary Table 5), that are widely known to be enriched for disease heritability [20–23]. We compared GCSC to MESC [10], stratified LD score regression [13] (S-LDSC) with the baseline-LD model [24, 25] (using S-LDSC to assess disease heritability enrichment of SNPs ±100kb from genes in a gene set, as previously described [3, 5]), and MAGMA [2]. Comparisons to MESC were restricted to the 22 diseases and complex traits analyzed in both this manuscript and ref. [10].

We first analyzed random gene sets with 10% of all genes in the gene set. Results of GCSC, MESC, S-LDSC and MAGMA for each disease/trait are reported in Supplementary Fig. 7. Although random sets of genes can be enriched or depleted for heritability, one would expect that these signals would be weak, and that random gene sets would be enriched or depleted for heritability with similar probabilities. Unexpectedly, we determined that S-LDSC gene set enrichment estimates for these random gene sets were biased, although S-LDSC regression coefficients (the primary focus of ref. [3]) were unbiased; we discuss possible reasons for the biased S-LDSC gene set enrichment estimates in the Supplementary Note, and focus on S-LDSC regression coefficients in our comparisons. GCSC, MESC and MAGMA produced unbiased results for random gene sets.

We next analyzed the two positive-control gene sets. Results of GCSC vs. MESC for each disease/trait are reported in Fig. 3A (high-pLI genes) and Fig. 3B (MGI essential genes). GCSC consistently attained higher power than MESC for both high-pLI genes (43/43 diseases/traits; regression of *−*log_10_ p-values: slope = 2.22) and MGI essential genes (40/43 diseases/traits; slope = 3.83). As in our simulations, we attribute the higher power of GCSC to the greater flexibility of GCSC. Results of GCSC vs. S-LDSC and GCSC vs. MAGMA for each disease/trait are reported in Supplementary Fig. 8. GCSC generally attained higher power than S-LDSC but lower power than MAGMA. However, MAGMA was previously reported to have an elevated type I error rate in simulations [3]. We also note that GCSC answers a different question than S-LDSC or MAGMA (pertaining specifically to disease heritability explained by predicted gene expression) and that GCSC outperforms MAGMA for some diseases/traits in the analyses below

**Figure 3:**
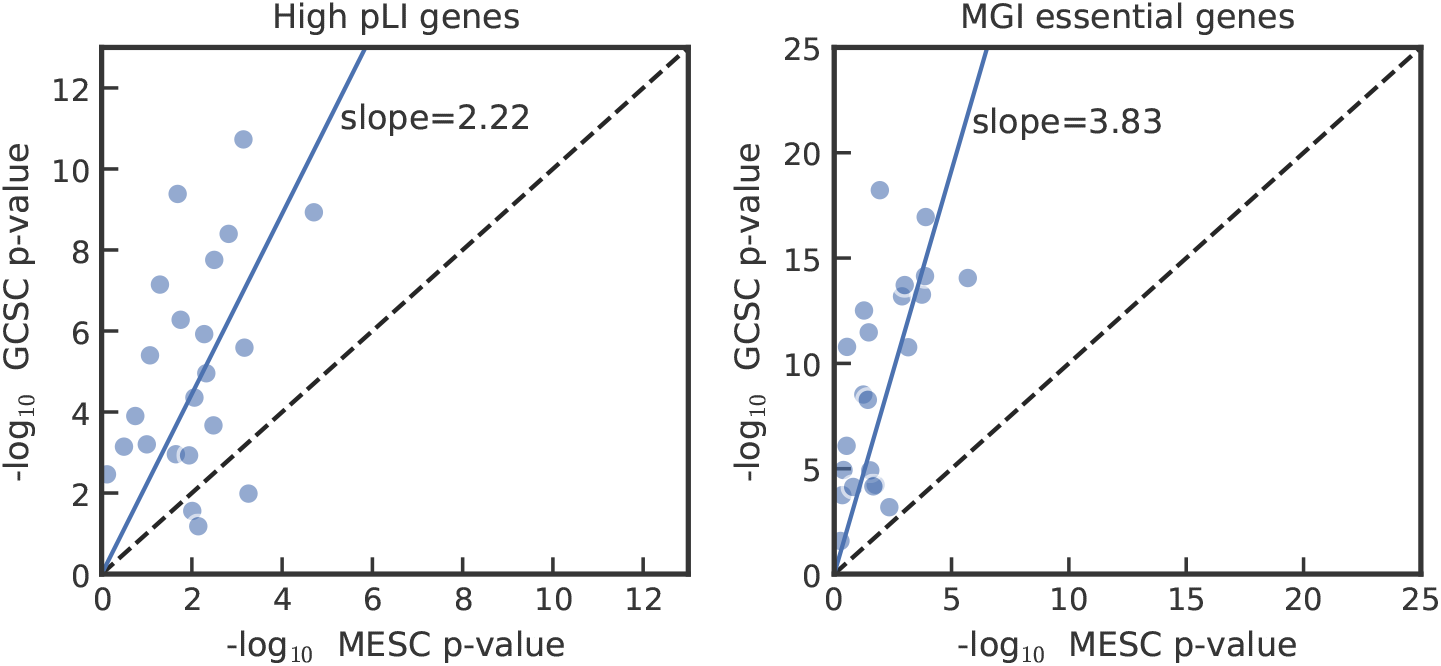
Performance of GCSC and MESC in analyses of positive-control gene sets. We report results for (A) high-pLI genes and (B) MGI essential genes. Analyses are restricted to the 22 diseases/traits analyzed in both this study and [10]. P-values are for enrichment relative to both protein-coding and non-protein-coding genes (see Methods). Numerical results are reported in Supplementary Table 5.

### Results for non-disease-specific gene sets

We applied GCSC to 38 non-disease-specific gene sets (Supplementary Table 5) and meta-analyzed the results across the 43 independent diseases and complex traits. Results are reported in Fig. 4 and Supplementary Table 6. Gene sets that were significantly enriched for disease heritability explained by predicted gene expression included several gene sets that are widely known to be enriched for disease heritability: ClinGen haploinsufficient genes, which are genes known to be pathogenic when their dosage is altered [26]; the high-pLI and MGI essential gene sets (also see Fig. 3); other constrained gene sets such as high-*s*_*het*_ [27]; and genes with high enhancer domain scores [28].

**Figure 4:**
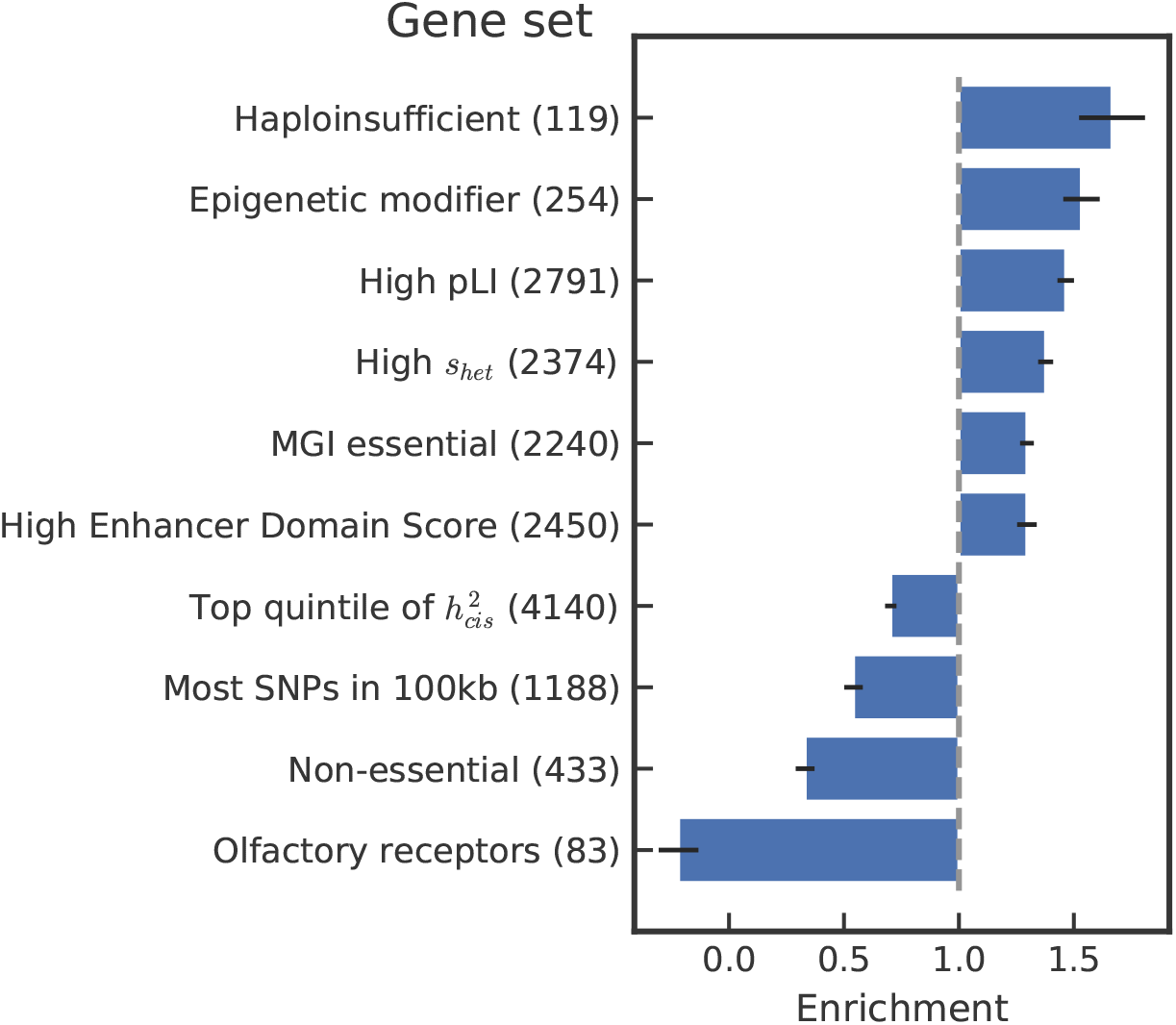
GCSC results for non-disease-specific gene sets. We report GCSC heritability enrichment estimates, meta-analyzed across 43 independent diseases/traits, for a subset of 10 non-disease-specific gene sets (numbers in parentheses denote the number of genes in the gene set with significant cis-heritability of gene expression in at least one tissue). Error bars denote 1 standard error. Numerical results, including results for all 38 non-disease-specific gene sets and all 43 independent diseases/traits, are reported in Supplementary Table 6.

Interestingly, epigenetic modifier genes [29] attained one of the largest enrichments. These genes have previously been shown to be enriched for heritability of neurological diseases [29] and cancer [30]. In contrast to these previous studies that analyzed a very limited set of diseases/traits, we determined that the enrichment of this gene set for heritability explained by predicted gene expression is ubiquitous across disease/traits, with enrichment greater than 1 for 39/43 diseases/traits analyzed (Supplementary Table 6).

In addition, we identified several gene sets that were depleted for disease heritability explained by predicted gene expression, in line with prior expectations. The strongest depletion was in olfactory receptors [31], which explained approximately zero heritability. Other depleted gene sets included genes with the most SNPs within 100kb, in line with negative selection purging genetic variation near important genes [32], and genes inferred to be non-essential in CRISPR assays [33].

We tested for evidence of 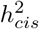 -dependent disease architectures. We determined that genes in the bottom quintile of 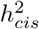 had 2.5 times higher disease heritability (*p* = 3.7 *×* 10^−17^) explained by predicted gene expression than genes in the top quintile of 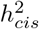 (Fig. 4 and Supplementary Table 7; this is unlikely to be an artifact of the method as it was not observed in simulations, Supplementary Fig. 9), consistent with previous studies reporting evidence that negative selection purges genetic variation impacting the expression of important genes [10, 34]. We repeated the analysis using quantile-normalized 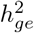 as a continuous-valued gene annotation (instead of 5 binary gene annotations) (see Methods), and detected an even stronger signal (*p*=1.6 *×* 10^−64^; Supplementary Table 8). In addition, we tested for evidence of co-regulation score-dependent disease architectures, but we did not find strong evidence that the disease heritability explained by a gene varies with its (all genes) co-regulation score (Supplementary Fig. 10 and 11).

### Results for biological pathways and specifically expressed gene sets

We applied GCSC to 11,394 biological pathways [5] (Supplementary Table 9) and 262 gene sets reflecting tissue-specific expression [3] (Supplementary Table 10) and analyzed each of the 43 diseases and complex traits separately, as enrichments for disease heritability explained by predicted gene expression for these gene sets are expected to be relatively disease-specific.

Results are reported in Fig. 5, Supplementary Fig. 12-16 and Supplementary Table 11. We identified 1,927 biological pathway-trait pairs and 412 specifically expressed gene set-trait pairs with enriched heritability explained by predicted gene expression (FDR<20%; Supplementary Tables 10 and 11). Many of the findings recapitulated known biology. Examples include vascular and cholesterol-related biological pathways and specifically expressed gene sets for total cholesterol (and other cardio-metabolic diseases/traits, e.g. coronary artery disease); immune-related biological pathways and specifically expressed gene sets for inflammatory bowel disease (and other autoimmune diseases, e.g. eczema); and growth-related biological pathways and specifically expressed gene sets for height.

**Figure 5:**
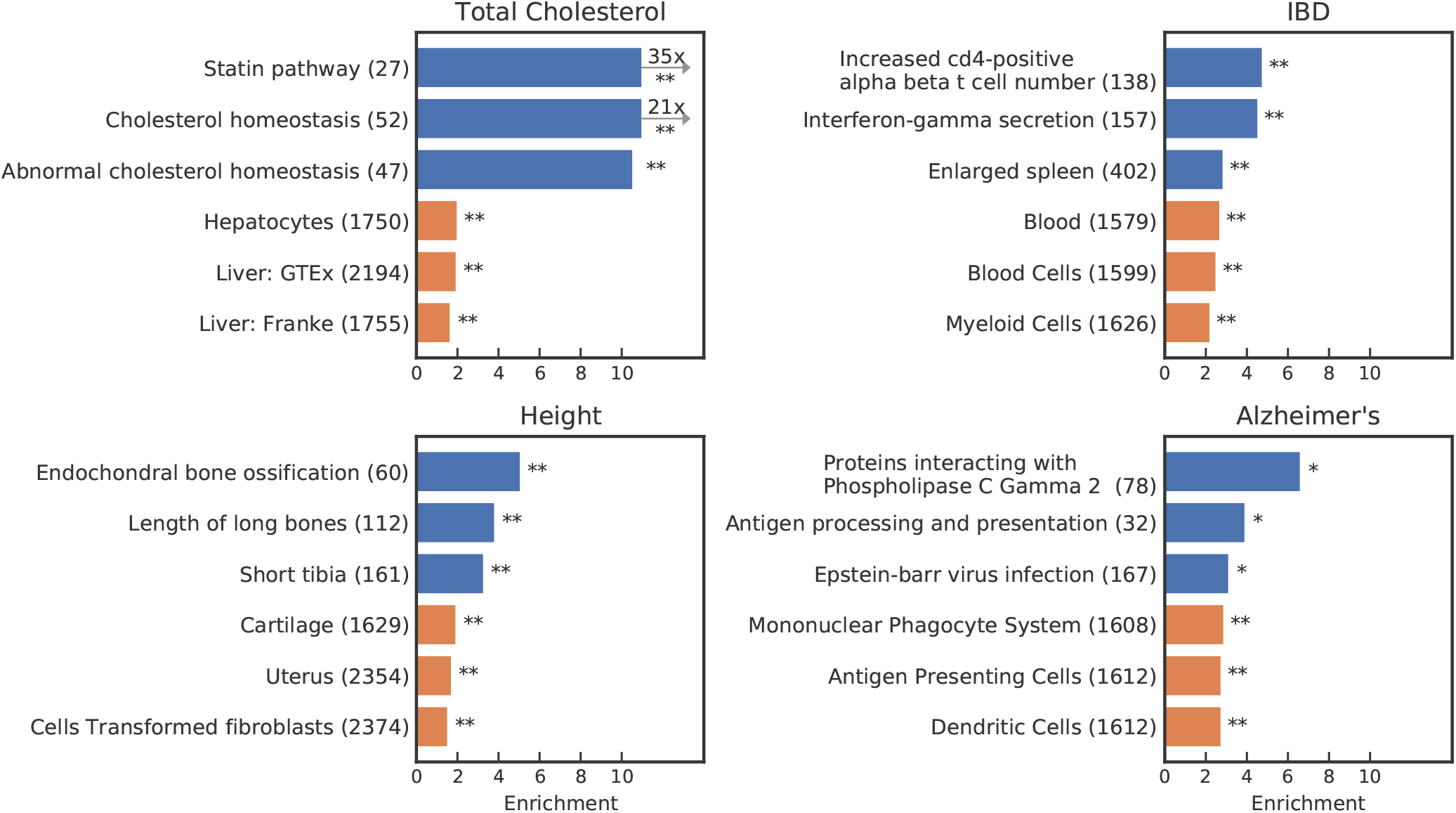
GCSC results for biological pathways and specifically expressed gene sets. We report GCSC heritability enrichment estimates for the top 3 most significantly enriched biological pathways (blue bars) and top 3 most significantly enriched specifically expressed gene sets (orange bars), for 4 diseases/traits: (A) total cholesterol, (B) inflammatory bowel disease (IBD), (C) height, and (D) Alzheimer’s disease (numbers in parentheses denote the number of genes in the gene set with significant cis-heritability of gene expression in at least one tissue). * denotes FDR<20%, ** denotes FDR<5%. Numerical results, including results for all 11,394 biological pathways and 262 specifically expressed gene sets for all 43 independent diseases/traits, are reported in Supplementary Table 11.

For Alzheimer’s disease, we identified evidence of an immune basis, and specifically a role for antigen presentation, in both top biological pathways and top specifically expressed gene sets enriched for heritability explained by predicted gene expression (Fig. 5). The most enriched Alzheimer’s disease gene set consisted of Phospholipase c gamma 2 (*PLCG2*) and its protein partners. Rare variants in *PLCG2* have been previously associated with late-onset Alzheimer’s disease, confirming the importance of this gene to Alzheimer’s disease [35]; a subsequent study showed that this gene plays a critical role in inflammation response in microglia [36]. This gene set contained 4 genes with transcriptome-wide significant TWAS associations (*P* < 0.05*/*15, 007 genes): *FCER1G* (Fc Fragment Of IgE Receptor Ig), *INPP5D* (Inositol Polyphosphate-5-Phosphatase D), *CD2AP* (CD2 Associated Protein) and *VASP* (Vasodilator Stimulated Phosphoprotein) (we note that *PLGC2* was not transcriptome-wide significant). The second most enriched Alzheimer’s disease gene set consisted of genes involved in antigen processing and presentation. Antigen presentation has been associated with Alzheimer’s disease [37, 38], however, a potential causal role remains unclear. This gene set contained no transcriptome-wide significant TWAS genes (*P* < 0.05*/*15, 007), indicating a more polygenic enrichment. Further supporting the role of the adaptive immune system in Alzheimer’s disease was a significant enrichment for genes involved in Epstein-Barr virus infection. This gene set contained two genes with transcriptome-wide significant TWAS associations (*P* < 0.05*/*15, 007 genes): *RELB* (RELB Proto-Oncogene, NF-KB Subunit) and *CR2* (Complement C3d Receptor 2). Some studies have reported evidence that prior Epstein-Barr virus infection may increase the risk of Alzheimer’s disease [39], but this remains disputed [40]. The 3 most significant specifically expressed cell types were all immune-related: mononuclear phagocyte system, antigen-presenting cells and dendritic cells. These three cell types are related (dendritic cells are strong antigen presenting cells and are a component of mononuclear phagoyctes), and have previously been associated with Alzheimer’s because of their role in clearing amyloid plaques [41, 42]. Overall, our results corroborate recent findings implicating the immune system as playing a major role in Alzheimer’s disease etiology [43, 44].

We note two other intriguing findings. First, two amyloid fiber formation gene sets were enriched for autism (Supplementary Fig. 14). Higher levels of amyloid-beta precursor have previously been linked to autism [45, 46], and the amyloid precursor protein secretase Cathepsin B *CTSB* was associated with autism in a previous TWAS [47]. Second, the gene set consisting of *ACTL6A* (Actin-like 6a) and its protein partners was enriched for educational attainment (Supplementary Fig. 12). *ACTL6A* was previously associated with intellectual disability [48] and is a part of the BAF complex, which is important for neurodevelopment [49]. However, to our knowledge, *ACTL6A* has not previously been associated with educational attainment. We also note two additional expected findings. First, a gene set consisting of genes involved in megakaryocyte development and platelet activation was enriched for platelet count (Supplementary Fig. 14). Second, two gene sets consisting of genes involved in circadian rhythms were enriched for chronotype (Supplementary Fig. 12).

We compared the results of GCSC to the results of MAGMA [2] for both non-disease-specific and disease-specific gene sets. For non-disease-specific gene sets and specifically expressed gene sets, GCSC and MAGMA had high concordance and a similar number of discoveries (Supplementary Fig. 17 and 18). For biological pathways, relative performance differed across diseases/traits. For several diseases/traits, including Alzheimer’s disease and autism, GCSC identified several significantly enriched gene sets that are consistent with known biology (see above and Supplementary Table 12), while MAGMA did not identify any significantly enriched gene sets (FDR< 20%). We did not compare to MESC and s-LDSC in these analyses because MAGMA generally attains higher power than MESC and S-LDSC in positive control gene sets, although MAGMA has been known to have an elevated type I error [3].

## Discussion

We developed a new method, GCSC, to identify gene sets that are enriched for disease heritability explained by the predicted expression of causal disease genes in the gene set. We verified via simulations that GCSC is well-calibrated, and well-powered to identify gene sets that are enriched for disease heritability explained by predicted expression. We determined that GCSC attains higher power than MESC [10] and S-LDSC [13], and identifies enriched gene sets that are complementary to those identified by MAGMA [2], including several significantly enriched gene sets for Alzheimer’s disease and autism that are consistent with known biology but are not identified by MAGMA. However, we note the GCSC estimand pertains to heritability *explained* by predicted expression (following the TWAS framework), in contrast to the MESC estimand which pertains to heritability *mediated* by predicted expression. In analyses of all genes, the more strict definition used by the MESC estimand addresses an important biological question [50].

Very recently, ref. [51] proposed a related method, GSR, that also identifies enriched gene sets by regressing TWAS *χ*^2^ statistics on gene co-regulation scores. However, we note 4 key differences between GCSC and GSR. First, GCSC uses a genomic block-jackknife to compute standard errors, attaining correct calibration (Supplementary Table 4); this is important, as analytical standard errors inherently assume independence between genes, an assumption that is violated by co-regulation between genes. Second, GCSC corrects for the bias arising from the distinction between predicted expression and the total cis-genetic component of expression; failure to apply this correction causes enrichment estimates to be biased (Supplementary Fig. 19). Third, GCSC uses a weighted regression to increase power. Fourth, our co-regulation score standardization procedure allows GCSC to be applied to multiple tissues to increase power.

We note several limitations of GCSC and its applications. First, GCSC restricts its analyses to genes with statistically significant cis-heritability of gene expression (analogous to TWAS). However, averaging across tissues, a majority (54%) of genes were included in our analyses, and this number will increase as gene expression data sets grow larger [52]. Second, our approach of concatenating data for multiple tissues to increase power does not account for differences between tissues; a potential solution is to explicitly model co-regulation across tissues (in addition to co-regulation across genes), which is an appealing direction for future research. Third, the choice of gene sets that we have analyzed is not comprehensive. In particular, analysis of specifically expressed gene sets derived from single-cell RNA-seq data [53] is a promising direction for future research. Fourth, GCSC may be impacted by pleiotropic effects involving SNPs that independently impact gene expression and disease risk (analogous to TWAS). However, GCSC pools information across different eQTLs for the same gene to leverage directional consistency, and pleiotropic effects are not expected to be directionally consistent. Finally, as with all analyses of enriched gene sets, an important caveat is that an enriched gene set representing a biological process may be enriched for reasons unrelated to the biological process, such that results should be interpreted cautiously. Despite these limitations, GCSC is a valuable tool for identifying enriched gene sets that provide insights about disease etiology.

## Code Availability

Software implementing GCSC will be made available at https://github.com/ksiewert prior to publication.

## Data Availability

All TWAS statistics and co-regulation scores are available for download at: https://alkesgroup.broadinstitute.org/GCSC/.

## Methods

### Gene expression models

Gene expression prediction models are a key input of the GCSC method (Fig. 1). A gene expression prediction model is a weighted combination of alleles that jointly predict the cis-genetic component of gene expression for a gene in a given tissue. We obtained these models from http://gusevlab.org/projects/fusion/ [6], which used GTEx (v7) [7]. For each tissue, we only retained gene expression prediction models for genes with significant (*p* < 0.05) 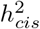, as estimated by GCTA [14].

### TWAS

We used FUSION with default parameters [6] to perform TWAS for real diseases/traits, and our own implementation of the FUSION test on simulated traits. The FUSION association statistic is:

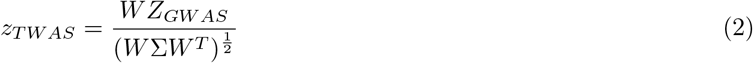

where *W* is the vector of eQTL weights from the gene’s expression model, and *Z*_*GW AS*_ is a vector of GWAS z-scores at each of the SNPs in the gene model.

### Calculation of co-regulation scores

To calculate co-regulation scores in each tissue, we first used the gene expression prediction models for that tissue (restricting to genes with significant (*p* < 0.05) 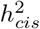) to predict gene expression in each of 503 1000 Genomes Europeans [15]. We then calculated the squared Pearson correlation between each pair of genes with gene bodies within 2 Mb of each other. The co-regulation score for each gene is the sum of these correlations across all nearby genes, including the gene itself.

However, the actual estimand involves the correlation between the *predicted* expression of the focal gene (since TWAS statistics are computed using predicted expression) and the *true cis-genetic component of* expression of a nearby gene (whose causal effect it tags) (see Supplementary Note). In the case of the correlation of a gene with itself, the correlation in predicted expression of the gene with itself (which is equal to 1) is an upwardly biased estimator of the correlation between the predicted expression of the gene and the true cis-genetic component of expression of the gene (which is less than 1, unless the gene expression prediction model is perfectly accurate). We therefore use a bias-corrected corrected estimate of a gene’s co-regulation with itself:

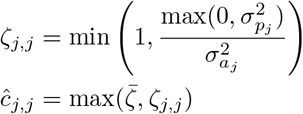

where 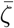 is the mean *ζ* across all genes in the given tissue, 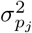 is the variance in gene expression explained by the gene expression model, which we estimate using the 5-fold cross validation accuracy of the gene model and 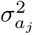 is the variance in gene expression explained by genetics. We used the GCTA estimate provided with the gene models for 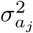. For a derivation of the bias of our estimator, see the Supplementary note. We take the minimum of the ratio of these two values and 1, because by random chance the ratio of the estimates of these two parameters can be above 1 even though the gene models cannot explain more variance than cis-genetics can explain. We find that this bias correction significantly reduced co-regulation score bias (Supplementary Fig. 20), while decreasing enrichment bias and maintaining power (Supplementary Fig. 19).

### GCSC regression

To run GCSC regression to assess enrichment of a gene set *S*, we regress TWAS *χ*^2^ statistics on two co-regulation scores for each gene *g*: *c*_*g*,0_, the sum of co-regulation scores of gene *g* with all nearby genes, and *c*_*g,S*_, corresponding to the sum of co-regulation scores of gene *g* with nearby genes in gene set *S*. The resulting slopes are proportional to *τ*_0_, the per-gene heritability explained by predicted expression of all genes without including the excess heritability contributed by genes in gene set *S* and *τ*_*S*_, the per-gene excess heritability contributed by genes in gene set *S*.

GCSC uses a weighted regression with three weights. The first is a tissue redundancy weight of 1*/n*_*tiss*_, where *n*_*tiss*_ is the number of tissues a gene has significant expression in (this weight is only applied in multi-tissue GCSC). This is to avoid weighting signal from genes found in many tissues more than more tissue-specific genes. Second, we include a co-regulation redundancy weight of 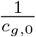 to avoid double-counting of signal from correlated genes; we note that this weight uses uncorrected co-regulation scores instead of the bias-corrected version. (This weight is analogous to the LD score weight of LD-score regression [54].) Third, we use a heteroscedasticity weight of 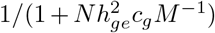, where M is the number of genes. We approximate 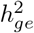 for use in the heteroscedasticity weight as 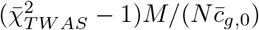, where 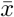 denotes the mean of x, analogous to the weight in [54]. Visualization of the linear relationship between TWAS *χ*^2^ and co-regulation scores in real data is available in Supplementary Fig. 21.

GCSC uses a free intercept to account for several possible scenarios that could drive the intercept away from 1: residual bias in co-regulation scores after correction, non-independence between co-regulation scores and gene-trait effect sizes, and the presence of SNP effects that are independent of co-regulation.

We use a weighted jackknife with 200 blocks of genes to calculate standard errors. To define these blocks, we divided the genome into 200 blocks of approximately equal number of genes. We define enrichment as:

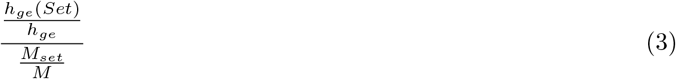

where *M* is the number of genes, *M*_*set*_ is the number of gene set genes. *h*_*ge*_ is the heritability explained by predicted gene expression, and *h*_*ge*_(*Set*) is the heritability explained by predicted gene expression (including effects of both *τ*_0_ and *τ*_*S*_) of gene set genes. Because enrichment is not normally distributed, we instead use the following normally distributed quantity to calculate a p-value, as done in [55]:

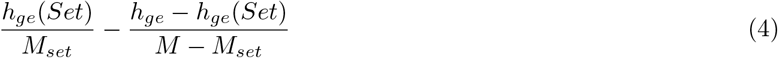

GCSC can be run using TWAS and co-regulation score pairs from a single tissue, or from a larger number of tissues concatenated. To correct for there being different numbers of cis-heritable genes in different tissues due to different sample sizes, we apply a correction to the co-regulation scores. This correction standardizes the “all genes” co-regulation scores in each tissue to have the same mean and variance as those in the tissue with the maximum number of significant genes. However, gene set co-regulation score standard deviations will be noisy, so we use a scaled version of the standard deviation of the all genes co-regulation score to standardize the gene set co-regulation score. Rescaled co-regulation scores, 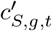 are:

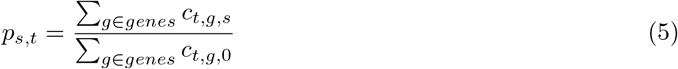

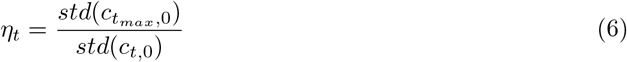

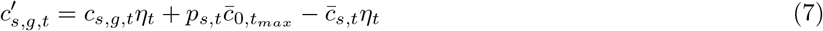

where *p*_*s,t*_ is the proportion of genes in tissue *t* in set *S, c*_*t,g,S*_ is the co-regulation score of gene *g* in tissue *t* for gene set *S* (*S* = 0 for the all genes score). *t*_*max*_ is the tissue with the highest mean co-regulation score, *g* is the total number of genes, std denotes standard deviation across genes and 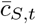 is the mean co-regulation score across genes for gene set *S* in tissue *t*.

We find that multi-tissue GCSC using all tissues outperforms single tissue GCSC in real data, even when the single tissue is the most biologically relevant tissue (Supplementary Fig. 22). We hypothesize this is due to the high correlation in cis-genetically determined gene expression between tissues, which was estimated to be 75% in [17]. In simulations, we find that single-tissue GCSC has similar power to multi-tissue GCSC when using the n=500 tissue, but substantially worse power for the n=80 (Supplementary Fig. 23). Due to the performance increase of multi-tissue GCSC in real data, we ran GCSC on all 48 tissues, except for our analyses of 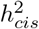 and co-regulation dependent architectures because we found that the lowest sample size tissue somewhat diluted the observed 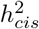 architecture from the expected in simulations.

In addition to estimating gene set enrichment, GCSC can be used to estimate the heritability explained by predicted gene expression, 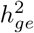, where 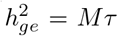, where M is the number of genes and *τ* is the per-gene heritability estimate from the univariable GCSC regression. This quantity includes heritability mediated by gene expression as well as heritability correlated with gene expression (for example, causal SNPs in linkage disequilibrium with eQTLs). Because 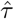 is defined as the mean effect across all genes in the regression, *M* should include only genes included in the regression. In single tissue GCSC, *M* is therefore the number of significantly heritable genes. In multi-tissue GCSC, there is no clear definition of *M*, as different tissues will have different number of significantly heritable genes. Therefore, when estimating 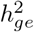 we use single-tissue GCSC. We find that in realistic simulations where heritability is split between two tissues (our default simulations), in the tissue with the higher eQTL sample sizes (n=500), single-tissue GCSC is only slightly downwardly biased (Supplementary Fig. 24). In contrast, in the smaller sample size (n=80), GCSC has a significantly downwardly biased estimate (Supplementary Fig. 24). This is likely due to the smaller number of significantly heritable genes that occurs, because GCTA standard errors increase as sample size decreases.

### Simulations

We performed extensive whole-genome simulations, which included using real genotypes from the UK Biobank [16] to simulate a GWAS with individuals with simulated gene expression and trait values, along with an eQTL study, a TWAS, and calculation of co-regulation scores. Our simulations were based on the TWAS simulator available at https://github.com/mancusolab/twas_sim [12]. We simulated expression in 10,000 genes which were randomly sampled from a list of all genes in the genome that contained at least 5 SNPs within 50kb from the gene edges, and the same 10,000 genes were selected for both tissues. Genes were randomly selected from among the 10,000 to be in the gene set.

We first simulated gene expression and resulting trait values for GWAS individuals. For each gene, we sampled the number of causal eQTLs from a uniform distribution between 1 and 5 and randomly selected the causal eQTLs from the set of SNPs within 50kb of the gene body. For ease of simulating correlated gene expression between tissues, the same eQTLs were chosen for the two tissues, and their effect sizes were drawn from a bivariable normal distribution with mean zero, variance specified to produce a total cis-heritability of gene expression in each tissue of 12%, and covariance specified to produce a cross-tissue cis-genetic correlation of 75% (following the estimate of [17]). For simulations with variable values of 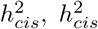 for each gene was drawn from an exponential distribution with parameter 0.12, and if the sampled 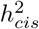 value was less than 0.01, it was raised to 0.01 to more closely mimic the distribution of 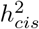 seen in real data. We restricted the set of SNPs used in the GWAS and gene expression simulations to those found in both 1000 Genomes European individuals and the UK Biobank individuals. We further restricted to SNPs that had a high info score in UK Biobank and had a minor allele frequency of at least 1%. In our default simulations, gene to trait effect sizes were randomly sampled to be positive or negative. In our simulations with variable gene to trait effect sizes, gene to trait effect sizes were drawn from a normal distribution with mean zero, a covariance between the tissues of zero, and a variance equal to that needed to get the desired 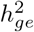. We verified that the resulting distributions of 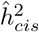 and co-regulation scores were similar to GTEx v7 (Supplementary Fig. 25 and 26).

We then simulated an eQTL study, where causal eQTL effects were used to impute expression into either 80 or 500 (non-overlapping) individuals, corresponding to reference panels from the two tissues, simulated from the LD matrix of 1000 Genomes, European individuals. Gene expression models were obtained using the genotypes and imputed expression values in the 80 and 500 individuals using LASSO with the regularization parameter determined using 5-fold cross-validation. In addition, 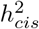 values for use in the bias correction and analysis of 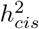 –dependent architectures were estimated using the GCTA package [14].

Simulated eQTL effects were then used to predict expression of each gene into each of the 50,000 GWAS individuals. We used individual-level data from British individuals in the UK Biobank (with relatives removed) [16] for the GWAS. We then performed a GWAS using a generalized linear model using plink [56], in which we used a default of 10% for the disease heritability explained by gene expression in each of the two tissues (summing to a total disease heritability of 20%-in line with real estimates of heritability mediated by gene expression by [10]).

We then calculated TWAS statistics and co-regulation scores. We first removed all simulated genes in each tissue with an insignificant (p> 0.05) GCTA estimate of 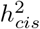, and performed a TWAS (see above). In addition, we imputed expression into 500 individuals simulated from the LD matrix of 1000 Genomes, European, and calculated “all genes” and “gene set” co-regulation scores.

We compared the distribution of p-values for false positive enrichments and depletions in null simulations. We found that GCSC is slightly conservative using a 1-sided test for enrichments, and anti-conservative using a 1-sided test for depletions, while the two-sided p-values appear well-calibrated (Supplementary Fig. 27).

We performed simulations with two types of 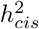 dependent architectures. Our default model was that 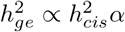, where *α* is the gene to trait effect size, in line with the model from [10]. In order to test the performance of GCSC under an alternative architecture, we performed simulations in which 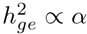 but independent of 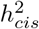. We found that GCSC captures both of these situations well (Supplementary Fig. 9 and 28).

### GWAS Summary Statistics

We constructed a list of 43 independent diseases and complex traits with publicly available GWAS summary statistics (squared genetic correlation [57] less than 0.1 (Supplementary Table 2), restricting to well-powered diseases/traits (z-score for nonzero SNP-heritability *≥* 6), similar to what was done in ref. [24]).

### Application to real gene sets

We performed two types of GCSC analyses: trait-specific and meta-analyzed across traits. In the latter, we applied GCSC to 43 independent traits and 38 gene sets that are not thought to have trait-specific relevance. We then performed a random-effects meta-analysis on the enrichment results for each gene sets across all independent traits. We obtained these gene sets from https://github.com/macarthur-lab/gene_lists and http://www.epigeneticmachinery.org.

To test for 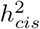 dependent architecture we assigned gene/tissue pairs into 5 bins of 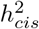 per tissue, then ran GCSC with 5 gene sets corresponding to the 5 quintiles. For co-regulation dependent architecture analyses, we divided co-regulation scores into ranges of size 0.5.

In order to generate more specific biological hypothesis, we also applied GCSC to a set of 11,394 gene sets from [5] that consisted of at least 20 genes and 262 sets of specifically expressed genes from [3]. These sets contained many gene sets with more narrow biological definitions than those that we meta-analyzed across traits, and thus we do not meta-analyze theses sets across traits, and instead look at trait-specific enrichment results.

We checked the distribution of enrichment p-values in the 11,394 gene sets from [5]. We found when applying two sided p-values, we observed an excess for low p-values, while when using one sided p-values testing for positive enrichment, we found a depletion (Supplementary Fig. 29 and 30), consistent to patterns seen in simulations. This observation indicates that GCSC finds that gene sets tend to be depleted, not enriched for heritability. This also indicates that using one sided p-values will result in a conservative test. For this reason, we perform two-sided tests, and filter for enrichment values greater than one. We used a Benjamini-Hochberg FDR procedure, and applied it separately to all specifically expressed gene sets and all 11,394 biological pathways for each trait.

### Application to MESC, s-LDSC and MAGMA

We applied s-LDSC using the baseline-LD model with no functional annotations, and mapped SNPs to genes using all SNPs within 100kb of each gene [5]. We used default settings in MESC [10]. For each comparison with MESC, GCSC was run on the set of all GTEx genes, instead of the set of only protein-coding GTEx genes, to match the background MESC uses. MAGMA v1.09 was run with no conditional analysis, and using a 10kb window around each gene [2].

## Acknowledgments

We thank Sasha Gusev, Luke O’Connor, Hilary Finucane, Omer Weissbrod, Tiffany Amariuta, Martin Zhang, Kushal Dey and Leandros Boukas for helpful discussions. This research was funded by NIH grants T32 DK110919, U01 HG009379, R01 MH101244, R37 MH107649, R01 MH115676 and R01 MH109978. This research was conducted using the UK Biobank Resource under Application 16549.

