## Supplementary Information for "Leveraging gene co-regulation to identify gene sets enriched for disease heritability"

#### CONTENTS

List of Figures 1

List of Tables 2

I. Supplementary Table Captions 3

II. Supplementary notes 4
- A. Derivation of GCSC regression equation 4
- B. Co-regulation score estimand and bias correction 6
- C. Possible reasons for biased S-LDSC gene set enrichment estimates for random gene sets 6
- D. Distinction between GCSC heritability estimand and MESC heritability estimand. 7

III. Supplementary Tables 8

IV. Supplementary Figures 10

#### I. SUPPLEMENTARY TABLE CAPTIONS

- **Supplementary Table 1. List of 48 GTEx tissues.** For each tissue, we report the GTEx (v7) sample size and number of protein-coding genes (of 18,094 total) with significant ($p < 0.05$) cis-heritability of gene expression, as estimated using GCTA.
- **Supplementary Table 2. List of 43 GWAS summary statistic data sets**. For each disease/trait, we report the disease/trait name, trait abbreviation, source, and sample size.
- **Supplementary Table 3. Numerical results of GCSC and MESC enrichment estimates**. We report enrichment estimates for each method in null (1x enrichment) and causal (3x enrichment) simulations, as well as the false positive rate and power at a p-value threshold of 5%.
- **Supplementary Table 4. Calibration of standard errors**. We report the mean standard error and the standard deviation of the mean for the gene set slope.
- **Supplementary Table 5. True positive gene set results**: MESC and GCSC results applied to high pLI genes and MGI non-essential genes.
- **Supplementary Table 6. Numerical GCSC results for non-disease-specific gene sets**. All genes (including non-protein coding genes) were included in the background set of genes in order to match the background set used by MESC. We report GCSC heritability enrichment estimates for each trait/gene pair as well as the meta-analysis results.
- **Supplementary Table 7. GCSC results of analyzing $h^2_{cis}$ quintile bins as gene sets to assess co-regulation dependent architectures**. We report the per gene heritability ($\tau$) explained by predicted gene expression of genes in quintiles of $h^2_{cis}$
- **Supplementary Table 8. GCSC results of analyzing quantile-normalized $h^2_{cis}$ as a continuous-valued gene annotation to assess co-regulation and $h^2_{cis}$ dependent architectures**. We report $h^2_{ge}$ and co-regulation dependent architecture in real data, meta-analyzed across 43 independent traits. In this analysis the effects of $h^2_{cis}$ and co-regulation were measure jointly in a single regression or were measured one at a time. $h^2_{cis}$ and co-regulation were quantile normalized across tissues, or not.
- **Supplementary Table 9. List of 11,394 biological pathways**. For each biological pathway, we list gene set IDs and descriptions for the gene sets. These gene sets are from Kim et al 2019 AJHG.
- **Supplementary Table 10. Numerical GCSC results for 262 specifically expressed gene sets**. For each specifically expressed gene set, we list the results of GCSC applied to 43 independent traits and 262 sets of specifically expressed genes.
- **Supplementary Table 11. Numerical GCSC results for biological pathways and specifically expressed gene sets**. Results of GCSC applied to 43 independent traits and biological pathways and specifically expressed gene sets.
- **Supplementary Table 12. Significantly enriched gene sets identified by GCSC that are consistent with known biology, but not identified by MAGMA** . We report the GCSC and MAGMA p-values for gene sets that pass a GCSC 20% FDR correction but not a MAGMA 20% FDR correction.

#### II. SUPPLEMENTARY NOTES

##### A. Derivation of GCSC regression equation

This subsection contains a derivation of Equation 1 (see Overview of methods), which states that the expected TWAS $\chi^2$ statistic for gene $g$ is

$$E(\chi^2_{TWAS_g}) = N\tau_0 c_{g,0} + N\tau_S c_{g,S} + 1, \tag{1}$$

where $N$ is GWAS sample size, $c_{g,0}$ is the co-regulation score of gene $g$ with respect to all genes (defined as $c_{g,0} = \sum_k r^2_{gk}$, where $r^2_{gk}$ is the squared correlation in predicted expression of gene $g$ and gene $k$), $c_{g,S}$ is the co-regulation score of gene $g$ with respect to genes in gene set $S$ (defined as $c_{g,S} = \sum_{k\in S} r^2_{gk}$), $\tau_0$ is the per-gene heritability explained by predicted expression of all genes without including the excess heritability contributed by genes in gene set $S$ and $\tau_S$ is the per-gene excess heritability contributed by genes in gene set $S$.

We start with a phenotypic model:

$$y = XU\alpha + X\beta + \epsilon \tag{2}$$

where $y$ is a length N vector of standardized phenotypes, $N$ is the number of individuals, $X$ is a $N \times s$ matrix of standardized genotypes, $s$ is the number of SNPs, $U$ is a $s \times M$ matrix of eQTL causal effects scaled so that $XU$ has mean 0 and variance 1, $\alpha$ is a length $M$ vector of gene to trait effect sizes, $\beta$ is a length $s$ vector of SNP effects (besides for eQTL effects) and $\epsilon$ is the error with mean zero. However, this model is not identifiable. We instead will be doing inference on the distribution of $\widetilde{\alpha}$, defined as the causal effect of gene expression on the trait plus any correlated effects of $\beta$. This avoids the identifiability issue of equation (1). Therefore:

$$\beta = U\gamma + \widetilde{\beta} \tag{3}$$

$$\widetilde{\alpha} = \alpha + \gamma \tag{4}$$

$$y = XU\widetilde{\alpha} + X\widetilde{\beta} + \epsilon \tag{5}$$

Where $\widetilde{\beta}$ is the component of $\beta$ that is uncorrelated with $W\alpha$ and $\gamma$ is the adjustment to $\alpha$ due to the presence of correlated effects to gene expression.

We define the marginal effect of gene $g$, $\hat{\alpha}_g$, as the following:

$$\hat{\alpha}_g = \frac{1}{N}(XW_g)^T y \tag{6}$$

$$= \frac{1}{N}(XW_g)^T(XU\widetilde{\alpha} + X\widetilde{\beta} + \epsilon) \tag{7}$$

$$= \frac{1}{N}\Big((XW_g)^T(XU\widetilde{\alpha}) + (XW_g)^T(X\widetilde{\beta}) + (XW_g)^T\epsilon\Big) \tag{8}$$

$$= \frac{1}{N}\big((XW_g)^T(XU\widetilde{\alpha}) + (XW_g)^T\epsilon\big) \tag{9}$$

where $W$ is a matrix of estimated causal eQTL effects generated using feature selection run on a gene expression panel (e.g. GTEx), standardized so that $XW$ has mean 0 and variance 1. We see in equation (9) that the marginal effect of gene $g$ depends on the covariance of gene $g$'s predicted expression and the true cis-genetically determined gene expression of each other gene. However, $U$ is unknown and must be approximated using $W$. We derive a bias correction for this approximation in section II B.

$$h^2_{ge} = Var[XW\widetilde{\alpha}] \tag{10}$$

$$= E_{U,\widetilde{\alpha}}[Var[XW\widetilde{\alpha}|X]] + Var_{W,\widetilde{\alpha}}[E[XW\widetilde{\alpha}|X]] \tag{11}$$

$$= E_{U,\widetilde{\alpha}}[\sum_{i\in Genes}(XW_i)^2\widetilde{\alpha}_i{}^2] \tag{12}$$

$$= ME[\widetilde{\alpha}^2] \tag{13}$$

$$E[\widetilde{\alpha}^2] = \frac{h^2_{ge}}{M} \tag{14}$$

where $h^2_{ge}$ is the variance in phenotype explained by predicted expression of eGenes.

We define the co-regulation score of gene $g$, $c_g$ as

$$\hat{r}_{g,k} = W_g^T X^T X W_k \tag{15}$$

$$\hat{c}_g = \sum_{i \in Genes} r^2(W_g X, W_i X) \tag{16}$$

where $r^2(W_g X, W_i X)$ is the squared correlation between imputed expression of gene $g$ and gene $i$.

After setting $U = W$ (which we will later correct for) we can therefore express the marginal gene to trait effect sizes as:

$$\hat{\alpha}_g = \sum_{k}^{genes} \hat{r}_{gk} \widetilde{\alpha}_k + \epsilon_g^{'} \tag{17}$$

Where $\epsilon_g^{'} = \frac{1}{N}(XW_g)^T \epsilon$ which has mean 0 and variance $\sigma_e^2/N$.

$$\chi^2_{TWAS_g} = N\hat{\alpha_g}^2 \tag{18}$$

$$E[\chi^2_{TWAS_g}] = E[N\hat{\alpha_g}^2] \tag{19}$$

$$= N\left(E\left[\left(\sum_{k \in genes} \hat{r}_{gk}\hat{\alpha}_k\right)^2\right]\right) \tag{20}$$

$$= N\left(\sum_{k \in genes} E[\hat{r}^2_{gk}]E\left[\widetilde{\alpha}^2_k\right] + E[\epsilon^2_g]\right) \tag{21}$$

$$= N\left(\sum_{k \in genes} \left(r^2_{gk} + \frac{1}{N}\right)\frac{h^2_{ge}}{M} + \frac{\sigma^2_e}{N}\right) \tag{22}$$

$$= N\hat{c}_g \frac{h^2_{ge}}{M} + \sigma^2_e + h^2_{ge} \tag{23}$$

$$= N\hat{c}_g \frac{h^2_{ge}}{M} + 1 \tag{24}$$

Since our phenotype is standardized to have variance 1, we replace $\sigma^2_e + h^2_{ge}$ with 1 to get from line 22 to 23. Note, we replace correlation with its expected values calculated using the formula of Wherry et al 1931 and assume co-regulation scores are independent of gene to trait effects in line 20. By regressing $\chi^2_{TWAS}$'s on $\hat{c}$'s, we therefore can estimate $h^2_{ge}$.

Next, we derive multivariable GCSC, in which one or more gene sets are being tested for enrichment. We start with a phenotypic model, where $\vec{\mathbb{1}}_{g \in s}$ is a vector of length M with 1 if gene $g$ is in the gene set $s$, and 0 otherwise. We define a gene set 0 as the set of all genes. Therefore, each $\widetilde{\alpha_s}$ represents additional heritability explained by the expression of genes in the set over that explained due to its presence in the set of all genes.

$$y = XU\widetilde{\alpha}_o + \sum_{s \in subsets} XU\vec{\mathbb{1}}_{g \in s}\widetilde{\alpha_s} + X\widetilde{\beta} + \epsilon \tag{25}$$

We define a partitioned co-regulation score $\hat{c}_g, s$ as the sum of squared correlation in predicted gene expression between gene g and genes in set $s$. We denote the set of all genes as set 0. Following similar steps to above, we obtain:

$$\chi^2_{TWAS_g} = N\Big(\hat{\alpha}_0 + \hat{\alpha}_S \vec{\mathbb{1}}_{g \in s}\Big)^2 \tag{26}$$

$$E[\chi^2_{TWAS_g}] = E\left[N\Big(\hat{\alpha}_0 + \hat{\alpha}_S \vec{\mathbb{1}}_{g \in s}\Big)^2\right] \tag{27}$$

$$= N\left(\hat{c}_{g,0}\frac{h^2_{ge,0}}{M} + \sum_{s \in sets} \hat{c}_{g,s}\frac{h^2_{ge,s}}{M}\right) + 1 \tag{28}$$

##### B. Co-regulation score estimand and bias correction

The actual co-regulation estimand involves the correlation between the *predicted* expression of the focal gene (since TWAS statistics are computed using predicted expression) and the *true cis-genetic component of* expression of a nearby gene (whose causal effect it tags) (see Supplementary Note). In other words, $E[\chi^2_{TWAS_j}]$ depends on the sum of the correlation between the predicted gene expression of gene $g$ and the actual gene expression of each other gene in GWAS individuals (denoted as $r^2(p_j, a_i)$). However, this correlation is unknown in reality, and instead must be estimated as the correlation between the predicted gene expression of gene $g$ and the predicted gene expression of each other gene in reference panel individuals $r^2(p_j, p_i)$. This leads to a slight upwards bias in the estimate of co-regulation of each gene with itself (when $i = j$). This is because $\hat{r}^2(p_j, p_j) = 1$, while $r^2(p_j, a_i)$ will usually be less than 1.

Let $a_i$ be the actual genetically determined gene expression of gene i, and define $p_i = a_i + \epsilon_i$. E$[a] = 0$ and E$[p] = 0$ We assume an infinitely large reference panel sample size (but not eQTL project sample size) to make the calculation tractable.

First, let $i = j$.

$$\text{Error}[r^2(p_i, p_j) \mid p_i, p_j] = r^2(p_i, p_j) - r^2(p_i, a_j) \tag{29}$$

$$= r^2(p_i, p_j) - \left(\frac{1}{\sigma_{p_i}\sigma_{a_j}}\text{Cov}(p_i, a_j)\right)^2 \tag{30}$$

$$= r^2(p_i, p_j) - \frac{1}{\sigma^2_{p_i}\sigma^2_{a_j}}(\text{Cov}(p_i, p_j) - \text{Cov}(p_i, \epsilon_j))^2 \tag{31}$$

$$= r^2(p_i, p_j) - \frac{1}{\sigma^2_{p_i}\sigma^2_{a_j}}(\text{Cov}(p_i, p_j) - \text{Cov}(a_i, \epsilon_j))^2 \tag{32}$$

$$= r^2(p_i, p_j) - \frac{\sigma^2_{p_j}}{\sigma^2_{p_j}\sigma^2_{p_i}\sigma^2_{a_j}}\text{Cov}^2(p_i, p_j) \tag{33}$$

$$= r^2(p_i, p_j) - \frac{\sigma^2_{p_j}}{\sigma^2_{a_j}} r^2(p_i, p_j) \tag{34}$$

Eq. (32) follows from equation (31) because $a_i$ is independent of $\epsilon_i$. We can then set $\sigma^2_{p_j}$ to be $r^2_{cv}$ (the 5-fold cross validation accuracy) and $\sigma^2_{a_j} = h^2_{exp}$ (The REML estimated cis heritability of gene expression) to give us the error of our estimator:

$$error = \hat{r}^2(p_i, p_j)\left(1 - \frac{r^2_{CV}}{h^2_{exp}}\right) \tag{35}$$

When $i \neq j$, the same derivation does not hold true, because $a_i$ may not be independent of $\epsilon_j$. However, we find that the co-regulation score estimates when $i \neq j$ are approximately unbiased without correction (see Supplementary Figure S20).

##### C. Possible reasons for biased S-LDSC gene set enrichment estimates for random gene sets

We determined that S-LDSC enrichment estimates for random gene sets (calculated with respect to the background set of genes, not all SNPs as is default) were biased, although S-LDSC regression coefficients (the primary focus of Finucane et al. 2018) were unbiased (Supplementary Fig. S7). We focus on S-LDSC regression coefficients in our comparisons between GCSC and S-LDSC, but discuss the bias in S-LDSC enrichment estimates here.

In detail, in analyses of real diseases/traits, S-LDSC enrichment estimates for random gene sets were biased in the sense that there was an excess of enrichment estimates >1 for some diseases/traits and an excess of enrichment esimates <1 for other traits (Supplementary Fig. S7). One possible cause for this could be violations to the linear model. The default procedure for running s-LDSC on gene sets is to have, among other annotations, two binary annotations: "SNPs near gene set genes" and "SNPs near all genes". Because the annotations are binary, a SNP near a single gene will have the same annotation value as a SNP near several genes. We hypothesize that this could result in a violation to the linear model, because SNPs near several genes might have a higher enrichment for heritability than

SNPs near a single gene. This would result in non-zero coefficients on the "SNPs near gene set genes" annotation, because like the "SNPs near all genes" annotation, it is correlated with the $\chi^2_{GWAS}$, but the model does not capture the true underlying relationship and the two annotations imperfectly capture the relationship in the data in different ways. Correcting this bias could be a direction of future work.

##### D. Distinction between GCSC heritability estimand and MESC heritability estimand.

GCSC estimates $h^2_{ge}$, the heritability explained by predicted gene expression, which includess effects both mediated by eQTLs and effects correlated with co-regulation scores. Because of this, $h^2_{ge}$ is expected to be larger than the mediated heritability, $h^2_{med}$, estimated by Yao et al. 2020. Yao et al. uses the baseline-LD model to control for pleiotropic effects of eQTLs, which could otherwise inflate estimates of $h^2_{med}$. In contrast, GCSC does not explicitly model pleiotropic effects because of its innate insensitivity to them. While MESC would otherwise be confounded by any non-eQTL effect in LD with an eQTL, GCSC estimates will only be affected if pleiotropic effects are directionally consistent with eQTL effects. An additional difference between the two methods is how they deal with data from multiple tissues. MESC combines gene expression models from all tissues into one gene model for each gene. In contrast, we find that for GCSC, it works better to instead use each gene/tissue pair as a unique point in the regression.

#### III. SUPPLEMENTARY TABLES

| Metric | GCSC | MESC |
|---|---|---|
| Enrichment, null sims | 1.004 (0.015) | 1.020 (0.022) |
| Enrichment, enriched sims | 2.784 (0.063) | 2.638 (0.092) |
| False positive rate | 0.026 (0.008) | 0.031 (0.008) |
| Power | 0.940 (0.022) | 0.567 (0.022) |

**Table S3:** We report enrichment estimates for each method in null (1x enrichment) and causal (3x enrichment) simulations, as well as the false positive rate and power at a p-value threshold of 5%, corresponding to figure 2.

| Proportion of genes in set | Mean standard error | Standard deviation of means |
|---|---|---|
| 0.005 | $1.1 \text{x} 10^{5}$ | $1.1 \text{x} 10^{5}$ |
| 0.01 | $7.3 \text{x} 10^{6}$ | $7.1 \text{x} 10^{6}$ |
| 0.05 | $3.4 \text{x} 10^{6}$ | $3.4 \text{x} 10^{6}$ |
| 0.1 | $2.5 \text{x} 10^{6}$ | $2.6 \text{x} 10^{6}$ |
| 0.2 | $1.9 \text{x} 10^{6}$ | $1.8 \text{x} 10^{6}$ |
| 0.4 | $1.5 \text{x} 10^{6}$ | $1.5 \text{x} 10^{6}$ |

**Table S4:** Calibration of gene set slope ($\tau_S$) standard errors in simulations with a 1x enrichment and with different proportions of genes in the gene set.

| $h^2_{cis}$ quintile | $\tau$ | standard error |
|---|---|---|
| 1 | 6.1e-6 | 3.8e-7 |
| 2 | 3.4e-6 | 2.7e-7 |
| 3 | 3.2e-6 | 2.5e-7 |
| 4 | 2.6e-6 | 2.2e-7 |
| 5 | 2.4e-6 | 2.2e-7 |

**Table S7:** $h^2_{ge}$ dependent architecture in real data, meta-analyzed across 43 independent traits.
We report the per gene heritability ($\tau$) explained by predicted gene expression of genes in quintiles of $h^2_{cis}$

| Joint | Type of architecture | Quantile normalized | $\tau*$ | P-value |
|---|---|---|---|---|
| No | $h^2_{cis}$ | False | -0.26 | $5.6 \text{x} 10^{-66}$ |
| No | Co-regulation | False | 0.09 | $4.4 \text{x} 10^{-7}$ |
| No | $h^2_{cis}$ | True | -0.26 | $1.6 \text{x} 10^{-64}$ |
| No | Co-regulation | True | 0.10 | $1.1 \text{x} 10^{-7}$ |
| Yes | $h^2_{cis}$ | False | -0.24 | $1.7 \text{x} 10^{-57}$ |
| Yes | Co-regulation | False | 0.06 | $3.5 \text{x} 10^{-4}$ |
| Yes | $h^2_{cis}$ | True | -0.24 | $2.3 \text{x} 10^{-55}$ |
| Yes | Co-regulation | True | 0.07 | $1.5 \text{x} 10^{-4}$ |

**Table S8:** $h^2_{ge}$ and co-regulation dependent architecture in real data, meta-analyzed across 43 independent traits. In this analysis the effects of $h^2_{cis}$ and co-regulation were measure jointly in a single regression or were measured one at a time. $h^2_{cis}$ and co-regulation were quantile normalized across tissues, or not. $\tau* = \frac{M sd_S \tau}{h^2_{ge}}$ where $sd_S$ is standard deviation of gene set.

| Trait | Gene set | GCSC p-value | MAGMA p-value |
|---|---|---|---|
| Alzheimer's | Phospholipase C gamma 2 | 1.68E-03 | 1.35E-02 |
| Alzheimer's | Antigen processing and presentation | 5.09E-03 | 5.91E-02 |
| Autism | Amyloid fiber formation | 1.93E-4 | 1.47E-01 |
| Blood pressure, Diastolic | Pericardial edema | 4.34E-03 | 5.31E-02 |
| Blood pressure, Diastolic | Myometrial Relaxation and Contraction Pathways | 5.72E-03 | 7.39E-02 |
| Blood pressure, Diastolic | Abnormal cardiovascular development | 1.26E-02 | 2.12E-02 |
| Bone mineral density, Heel tscore | Abnormal vertebral body morphology | 1.22E-02 | 3.30E-02 |
| Bone mineral density, Heel tscore | Abnormal zygomatic bone morphology | 1.28E-02 | 3.52E-02 |
| Cholesterol | Low density lipoprotein receptor | 1.66E-02 | 1.67E-01 |
| Cholesterol | Apolipoprotein E | 3.24E-02 | 3.41E-02 |
| Eczema | Increased interleukin-2 secretion | 3.97E-03 | 2.20E-02 |
| Eczema | Keratin 7 | 4.01E-03 | 3.78E-01 |
| Eczema | Major histocompatibility complex, class II, DR beta 3 | 8.23E-03 | 1.34E-02 |
| Eczema | Keratin 6B | 1.01E-02 | 3.10E-01 |
| Eczema | Keratin 6A | 1.02E-02 | 3.20E-01 |
| Eczema | Keratin 75 | 1.20E-02 | 5.69E-02 |
| Eczema | MHC class II antigen presentation | 1.35E-02 | 1.90E-01 |
| Eczema | Keratin 1 | 1.36E-02 | 2.75E-01 |
| Eczema | Keratin 6C | 1.49E-02 | 5.74E-02 |
| Eczema | Decreased IgG3 level | 1.52E-02 | 1.82E-02 |
| Educational Attainment | Actin like 6A | 9.25E-04 | 3.11E-03 |
| Height | Growth hormone receptor | 2.39E-03 | 2.31E-02 |
| Height | Growth hormone 1 | 3.34E-03 | 2.49E-01 |
| Height | Cell cycle | 5.05E-03 | 3.49E-02 |
| Height | Cell division cycle 20 | 5.75E-03 | 6.11E-01 |
| Height | Abnormal embryo size | 1.27E-02 | 3.47E-02 |
| IBD | TNFs binding their physiological receptors | 1.76E-03 | 4.68E-02 |
| IBD | TNFR2 non-canonical NF-kB pathway | 5.78E-03 | 8.91E-02 |
| Lung FVCzSMOKE | Abnormal lung morphology | 1.14E-02 | 2.25E-02 |
| Morning person | Shortened circadian behavior period | 8.46E-04 | 1.57E-03 |
| Morning person | Period circadian regulator 2 | 3.62E-03 | 3.91E-03 |
| Morning person | Melatonin metabolism and effects | 7.95E-03 | 2.02E-02 |
| Neuroticism | Synapse | 6.31E-04 | 1.80E-03 |
| Neuroticism | Reduced long term potentiation | 1.06E-02 | 2.02E-02 |
| Neuroticism | Dendrite | 1.34E-02 | 2.27E-02 |
| Neuroticism | Impaired contextual conditioning behavior | 1.34E-02 | 2.01E-02 |
| Neuroticism | Abnormal brain commissure morphology | 1.40E-02 | 1.62E-02 |
| Reaction Time | Abnormal excitatory postsynaptic potential | 1.34E-03 | 6.96E-03 |
| Testosterone, Male | Constitutive Androstane Receptor Pathway | 1.59E-02 | 3.06E-02 |
| White blood cell count | Chronic inflammation | 1.36E-02 | 4.51E-02 |

**Table S11:** Selection of gene sets that pass a GCSC 20% FDR correction but not a MAGMA 20% FDR correction.

### IV. SUPPLEMENTARY FIGURES

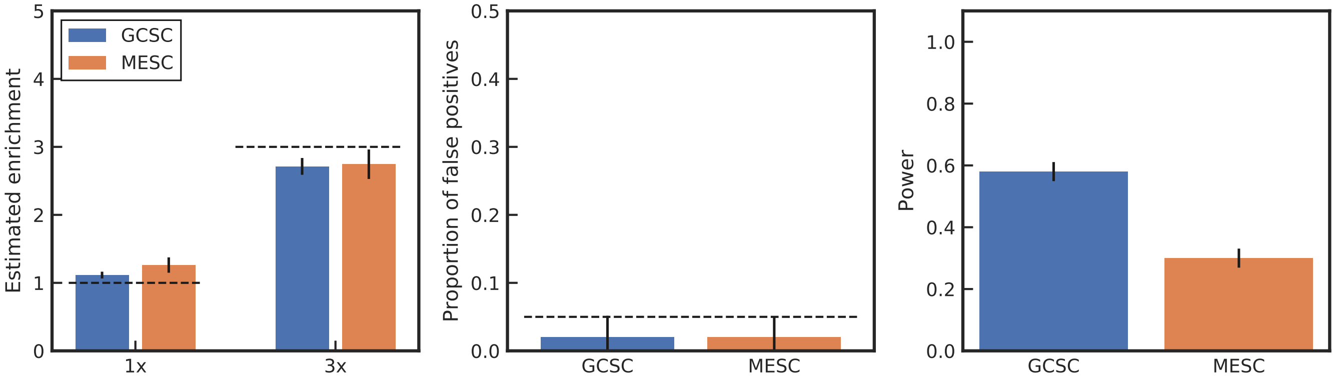

**Figure S1: Performance of GCSC and MESC in simulations in which the cis-heritability of gene expression and the disease explained by gene expression vary across genes.** Performance of GCSC and MESC on simulated data in which gene to trait effect sizes and $h^2_{cis}$ values are drawn from distributions instead of being constant. 10,000 genes were simulated, of which a random 20% were in the gene set. The GWAS sample size of 50,000 was used. Two tissues were simulated, one with a sample size of 80, and the other with 500. Both tissues had a fixed $h^2_{ge} = .12$ and each tissue explained 10% of trait variance. To model sharing of eQTLs between tissues, eQTL effect sizes has a correlation of 75% between tissues. In Panel (A), the estimated enrichment is plotted in simulations where there is no heritability enrichment in the gene set (1x) or when there is a 1.3x enrichment in the gene set (1.3x). In Panel (B), the proportion of 1x simulations in which an enrichment p-values less than 0.05 is shown. In Panel (C) the proportion of 3x simulations that each method reported an enrichment p-value less than 0.05 is shown. Error bars represent 1 standard error.

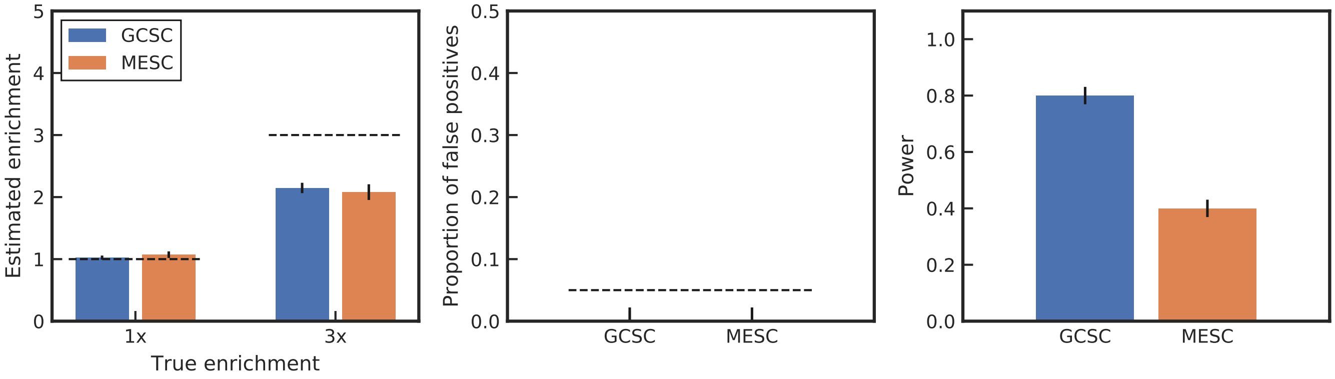

**Figure S2: Performance of GCSC and MESC in simulations with SNP effects independent of eQTL effects.** Performance of GCSC and MESC on simulated data, when half of heritability is not mediated by gene expression. The GWAS sample size of 50,000 was used. Two tissues were simulated, one with a sample size of 80, and the other with 500. Both tissues had a fixed $h^2_{ge} = .12$ and each tissue explained 10% of trait variance, with an additional 20% explained by non-eQTL effects of SNPs. To model sharing of eQTLs between tissues, eQTL effect sizes has a correlation of 75% between tissues. In Panel (A), the estimated enrichment is plotted in simulations where there is no heritability enrichment in the gene set (1x) or when there is a 1.3x enrichment in the gene set (1.3x). In Panel (B), the proportion of 1x simulations in which an enrichment p-values less than 0.05 is shown. In Panel (C) the proportion of 3x simulations that each method reported an enrichment p-value less than 0.05 is shown. Error bars represent 1 standard error.

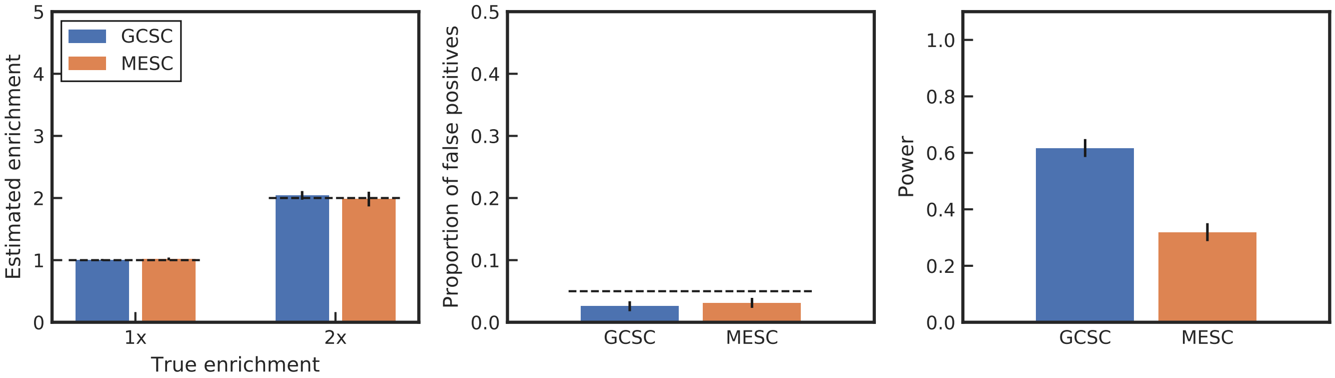

**Figure S3: Performance of GCSC and MESC in simulations with 2x enrichment instead of 3x enrichment.** Performance of GCSC and MESC on simulated data. The GWAS sample size of 50,000 was used. Two tissues were simulated, one with a sample size of 80, and the other with 500. Both tissues had a fixed $h^2_{ge} = .12$ and each tissue explained 10% of trait variance, with an additional 20% explained by non-eQTL effects of SNPs. To model sharing of eQTLs between tissues, eQTL effect sizes has a correlation of 75% between tissues. 1% of genes were in gene set. In Panel (A), the estimated enrichment is plotted in simulations where there is no heritability enrichment in the gene set (2x) or when there is a 2x enrichment in the gene set (2x). In Panel (B), the proportion of 1x simulations in which an enrichment p-values less than 0.05 is shown. In Panel (C) the proportion of 3x simulations that each method reported an enrichment p-value less than 0.05 is shown. Error bars represent 1 standard error.

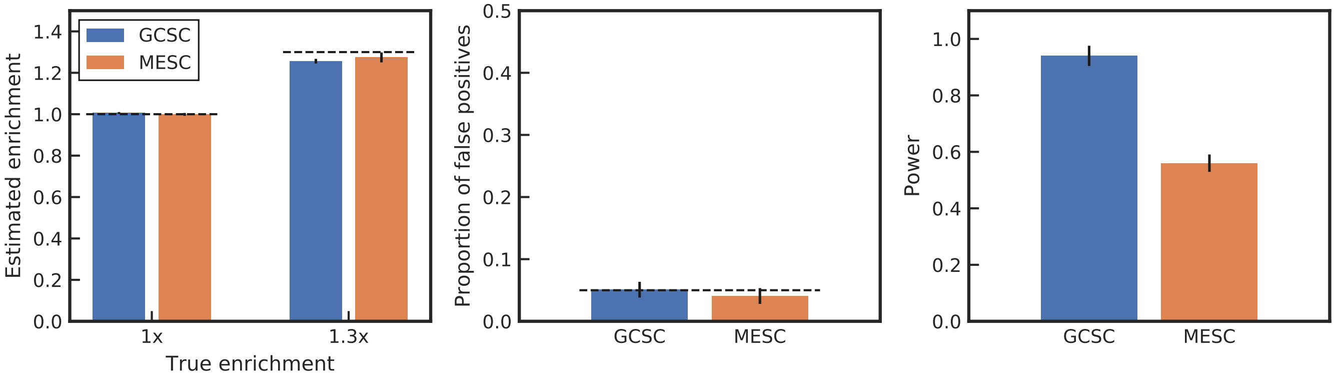

**Figure S4: Performance of GCSC and MESC in simulations with larger gene set.** Performance of GCSC and MESC on simulated data. 10,000 genes were simulated, of which a random 20% were in the gene set. The GWAS sample size of 50,000 was used. Two tissues were simulated, one with a sample size of 80, and the other with 500. Both tissues had a fixed $h^2_{ge} = .12$ and each tissue explained 10% of trait variance. To model sharing of eQTLs between tissues, eQTL effect sizes has a correlation of 75% between tissues. In Panel (A), the estimated enrichment is plotted in simulations where there is no heritability enrichment in the gene set (1x) or when there is a 1.3x enrichment in the gene set (1.3x). In Panel (B), the proportion of 1x simulations in which an enrichment p-values less than 0.05 is shown. In Panel (C) the proportion of 1.3x simulations that each method reported an enrichment p-value less than 0.05 is shown. Error bars represent 1 standard error.

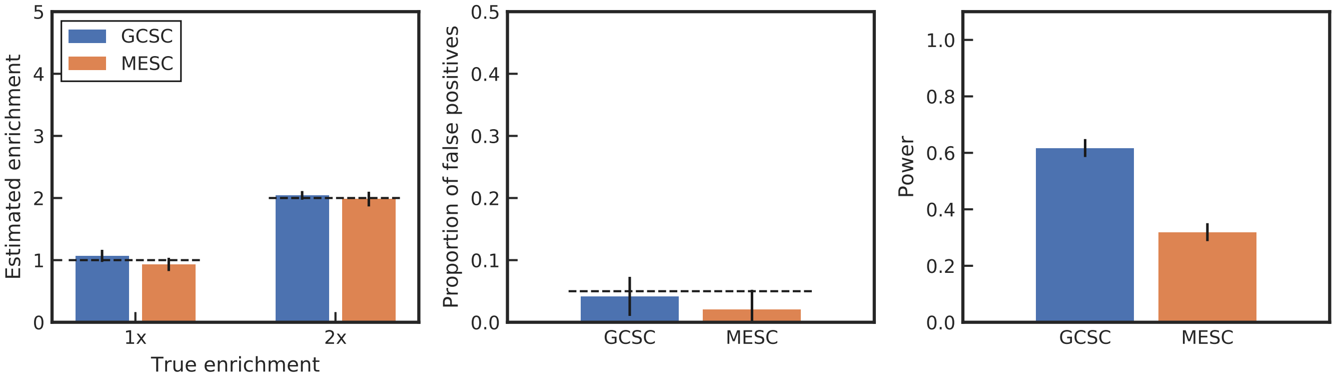

**Figure S5: Performance of GCSC and MESC in simulations with lower $h^2_{ge}$.** Performance of GCSC and MESC on simulated data. The GWAS sample size of 50,000 was used. Two tissues were simulated, one with a sample size of 80, and the other with 500. Both tissues had a fixed $h^2_{ge} = .12$ and each tissue explained 5% of trait variance. To model sharing of eQTLs between tissues, eQTL effect sizes has a correlation of 75% between tissues. 1% of genes were in gene set. In Panel (A), the estimated enrichment is plotted in simulations where there is no heritability enrichment in the gene set (2x) or when there is a 2x enrichment in the gene set (2x). In Panel (B), the proportion of 1x simulations in which an enrichment p-values less than 0.05 is shown. In Panel (C) the proportion of 3x simulations that each method reported an enrichment p-value less than 0.05 is shown. Error bars represent 1 standard error.

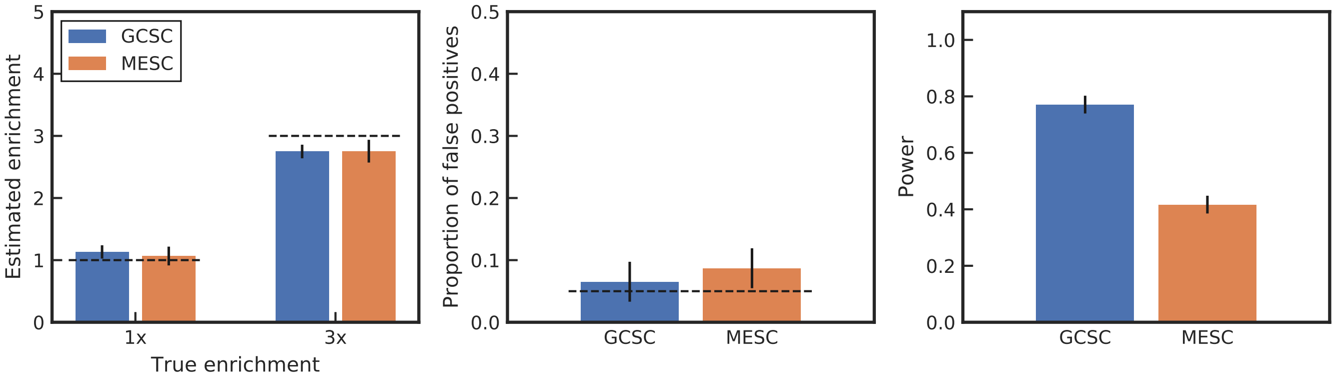

**Figure S6: Performance of GCSC and MESC in simulations with GWAS sample size reduced to 20,000.** Performance of GCSC and MESC on simulated data with 20,000 GWAS individuals. 10,000 genes were simulated, of which a random 1% were in the gene set. Two tissues were simulated, one with a sample size of 80, and the other with 500. Both tissues had a fixed $h^2_{ge} = .12$ and each tissue explained 10% of trait variance. To model sharing of eQTLs between tissues, eQTL effect sizes had a correlation of 75% between tissues. Tests are one-sided tests for enrichment. In Panel (A), the estimated enrichment is plotted in simulations where there is no heritability enrichment in the gene set (1x) or when there is a 3x enrichment in the gene set (3x). In Panel (B), the proportion of 1x simulations in which an enrichment p-values less than 0.05 is shown. In Panel (C) the proportion of 3x simulations in which each method reported an enrichment p-value less than 0.05 is shown. Error bars represent 1 standard error.

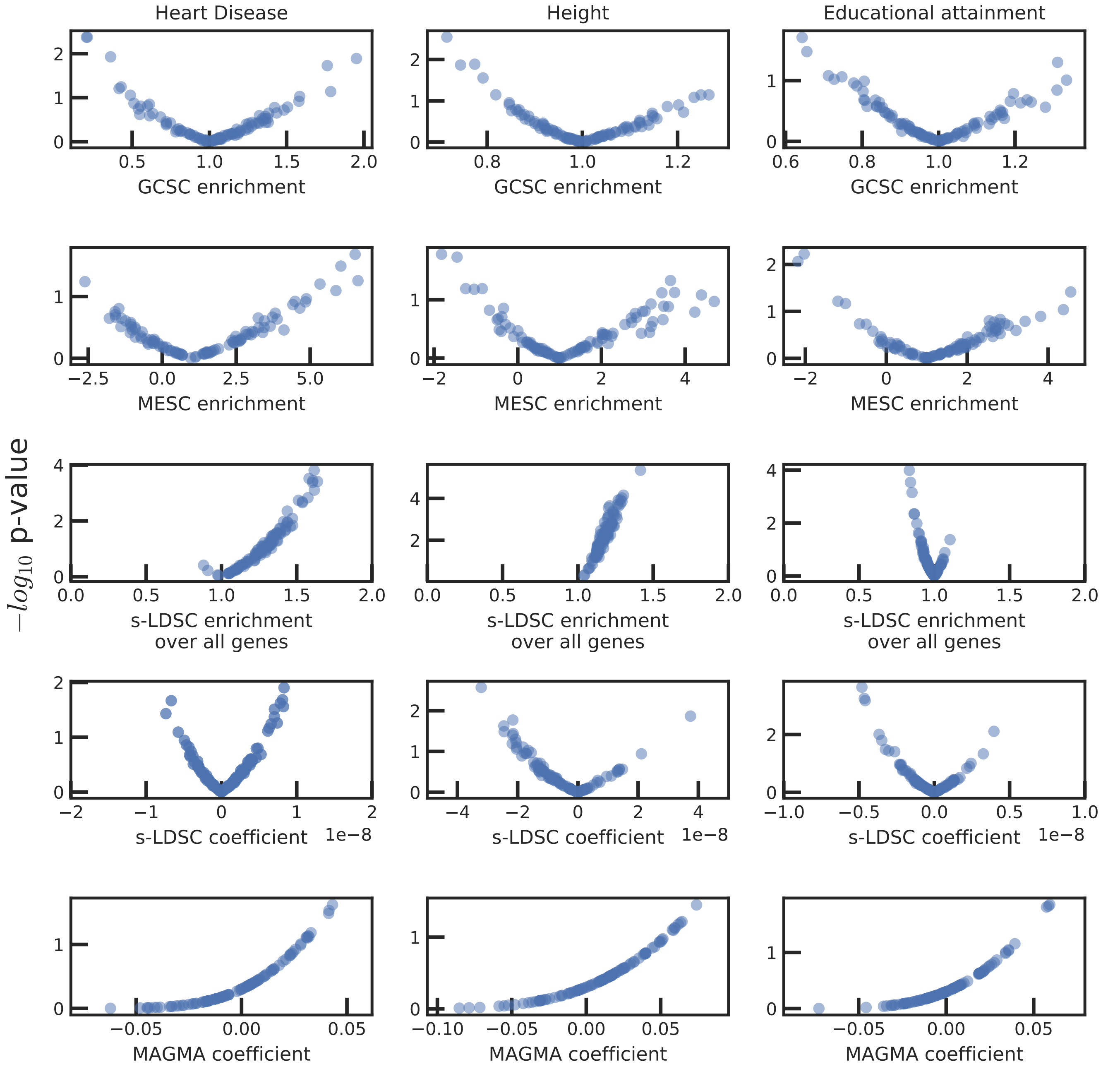

**Figure S7: Results of GCSC, MESC, S-LDSC and MAGMA in analyses of real data using random gene sets.** Enrichment estimates or coefficients and p-values when each method is applied to 100 random gene sets of size 10% of genes and 3 different traits. MAGMA performs a 1-sided test by default, while a 2-sided test was used for the other 3 methods.

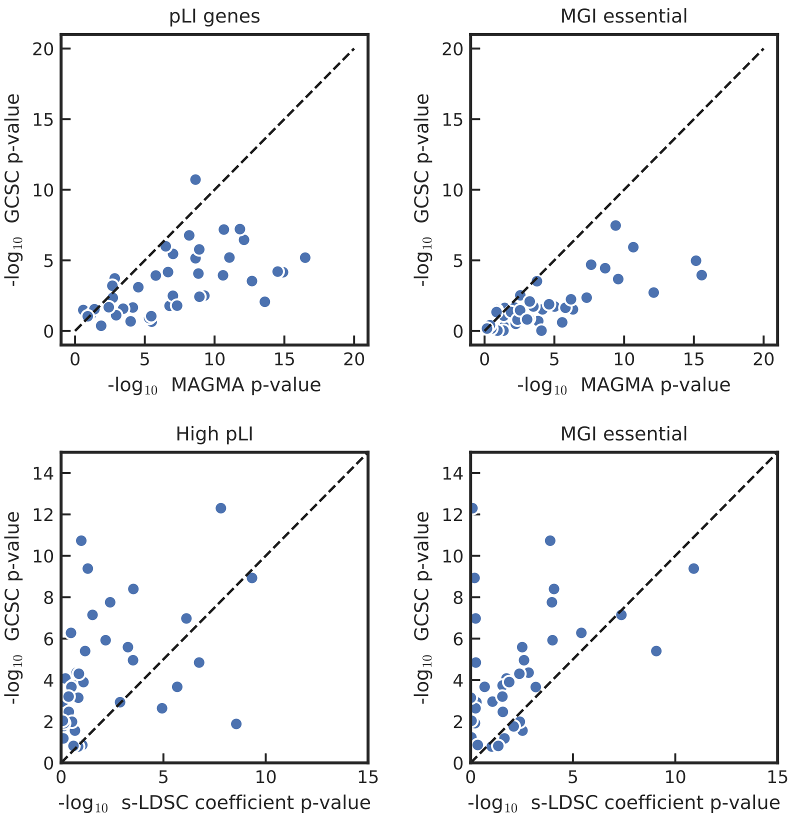

**Figure S8: Results of GCSC vs. S-LDSC and GCSC vs. MAGMA in analyses of real data using positive-control gene sets.** Performance on high pLI and MGI essential genes for MAGMA versus GCSC (top row) and s-LDSC versus GCSC (bottom) row. For the s-LDSC versus GCSC comparison, the background set of genes was all genes in order to compare to MESC. In the top row, only protein coding genes were used to match the background set of genes used in MAGMA.

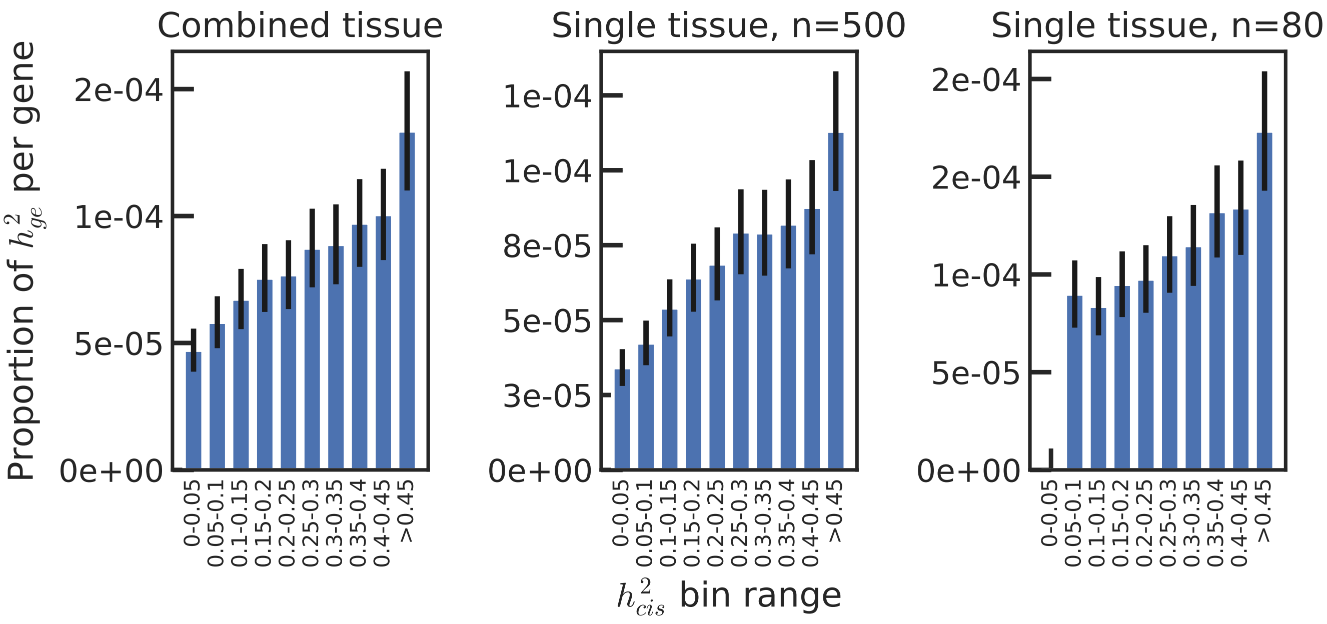

**Figure S9: No artifactual signals of $h^2_{cis}$-dependent architectures in simulations.** Per-gene heritability explained by predicted gene expression in simulations, by $h^2_{cis}$. Two tissues of sample sizes 500 and 80 were simulated and GCSC was performed using both tissues, or just one. Error bars denote +/- one standard error. Both $h^2_{cis}$ and gene to trait effect sizes were drawn from a distribution (see Methods).

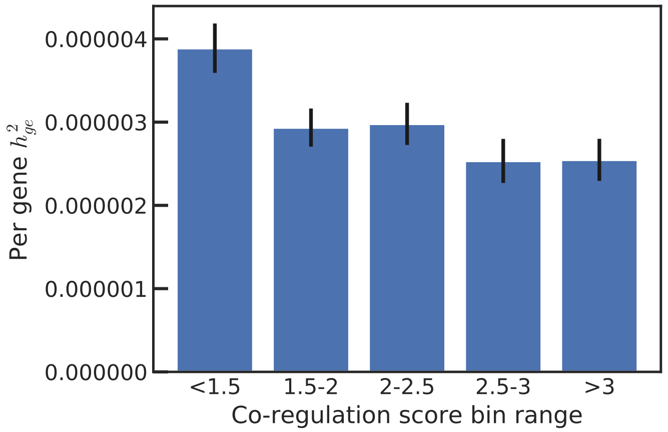

**Figure S10: GCSC results on co-regulation dependent architectures in analyses of real data.** Meta-analysis across traits of the proportion of the per gene heritability by co-regulation score bin. Per gene heritability was meta-analyzed across 43 independent traits. Error bars are +- 1 standard error.

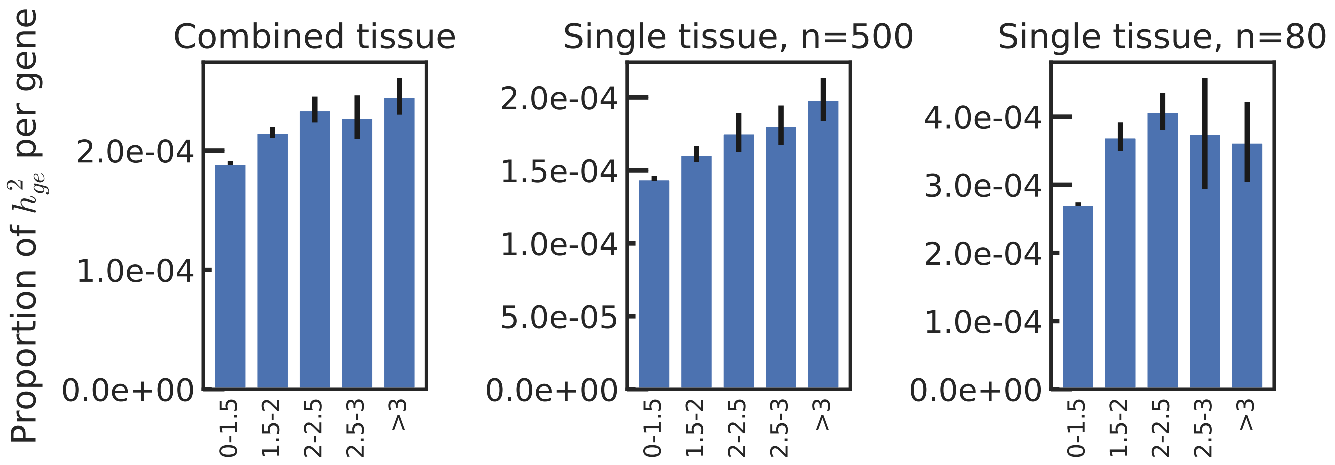

**Figure S11: GCSC results on co-regulation dependent architectures in simulations.** Per-gene heritability explained by predicted gene expression in simulations, by co-regulation score. Two tissues of sample sizes 500 and 80 were simulated and GCSC was performed using both tissues, or just one. Error bars denote +/- one standard error. Both $h^2_{cis}$ and gene to trait effect sizes were drawn from a distribution (see Methods).

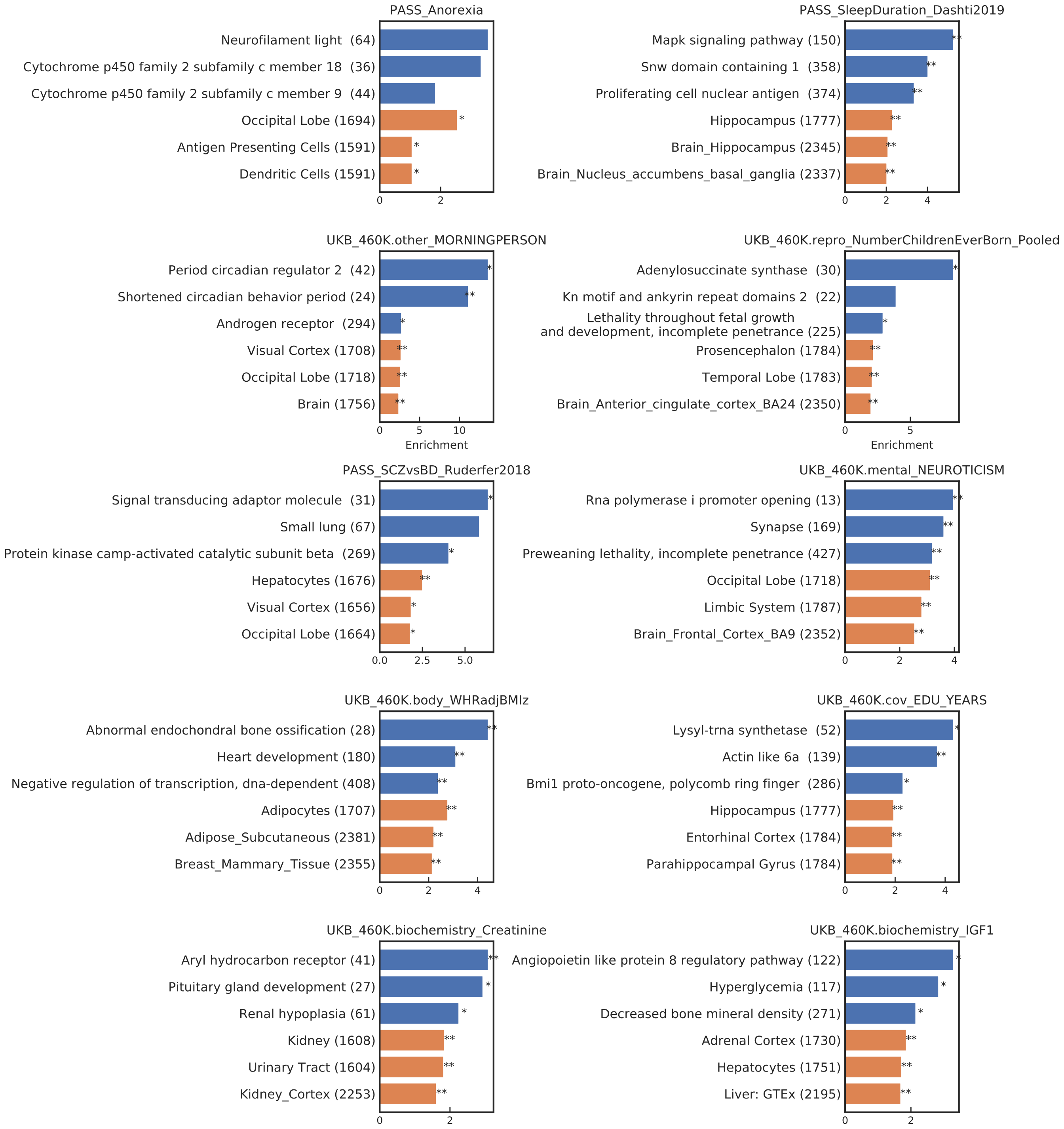

**Figure S12: Numerical GCSC results for biological pathways and specifically expressed gene sets for 10 of 43 diseases/traits.** Enrichment of gene sets in 4 traits with respect to the background of all protein coding genes. The top 3 most significant enrichment values from the specifically expressed gene analysis (in orange) and from the single trait gene set enrichment analysis (in blue) are displayed here, sorted by enrichment value. Error bars denote plus/minus one standard error. Number in parenthesis is the number of genes with a significant heritability of gene expression in at least one tissue. ** indicates passing the 20% FDR threshold and * indicates passing the 5% threshold. Gene sets pathways with the same descriptor are from different source (see supplementary table X).

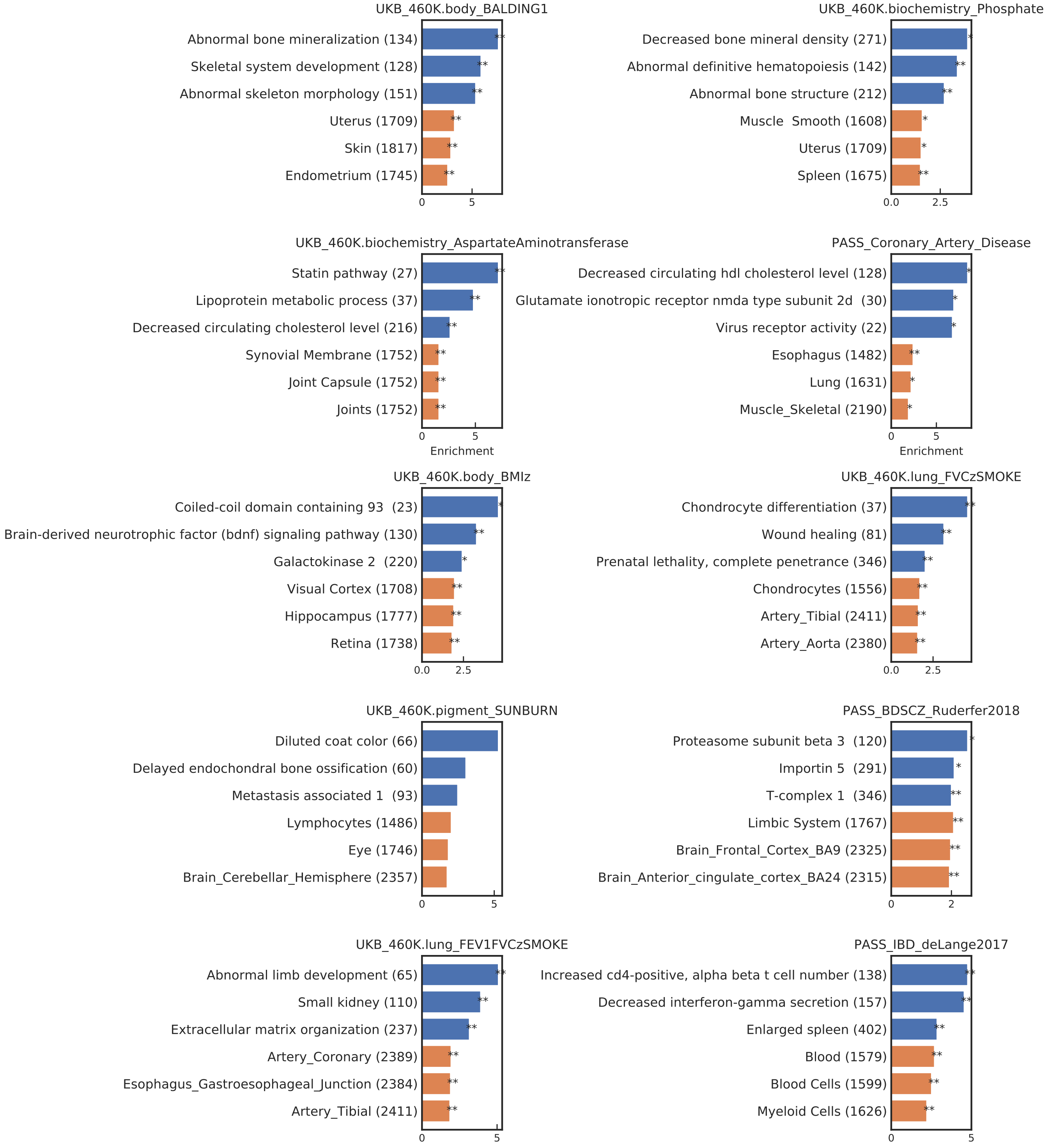

**Figure S13: Numerical GCSC results for biological pathways and specifically expressed gene sets for 10 of 43 diseases/traits.** Enrichment of gene sets in 4 traits with respect to the background of all protein coding genes. The top 3 most significant enrichment values from the specifically expressed gene analysis (in orange) and from the single trait gene set enrichment analysis (in blue) are displayed here, sorted by enrichment value. Error bars denote plus/minus one standard error. Number in parenthesis is the number of genes with a significant heritability of gene expression in at least one tissue. ** indicates passing the 20% FDR threshold and * indicates passing the 5% threshold. Gene sets pathways with the same descriptor are from different source (see supplementary table X).

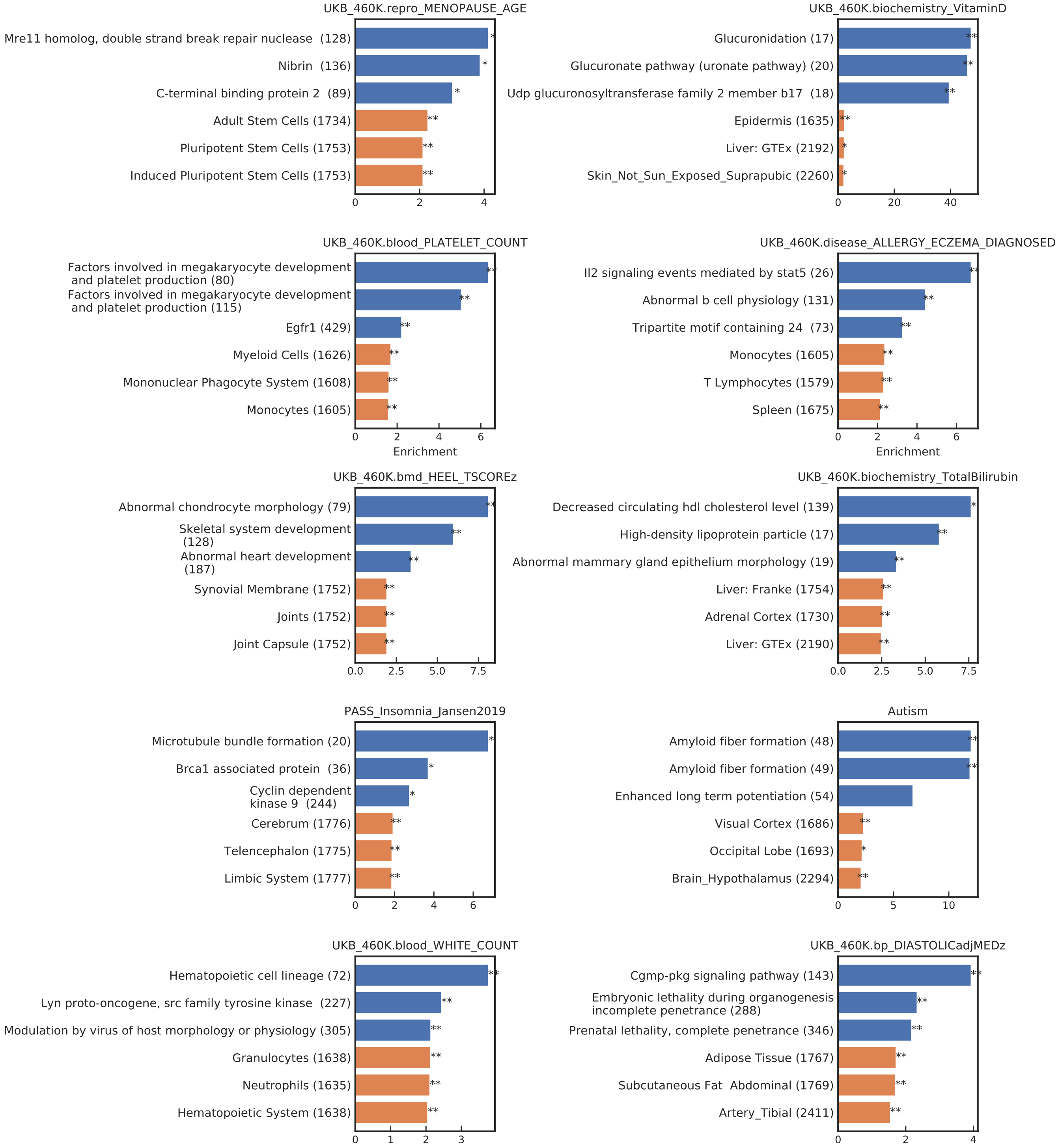

**Figure S14: Numerical GCSC results for biological pathways and specifically expressed gene sets for 10 of 43 diseases/traits.** Enrichment of gene sets in 4 traits with respect to the background of all protein coding genes. The top 3 most significant enrichment values from the specifically expressed gene analysis (in orange) and from the single trait gene set enrichment analysis (in blue) are displayed here, sorted by enrichment value. Error bars denote plus/minus one standard error. Number in parenthesis is the number of genes with a significant heritability of gene expression in at least one tissue. ** indicates passing the 20% FDR threshold and * indicates passing the 5% threshold. Gene sets pathways with the same descriptor are from different source (see supplementary table X).

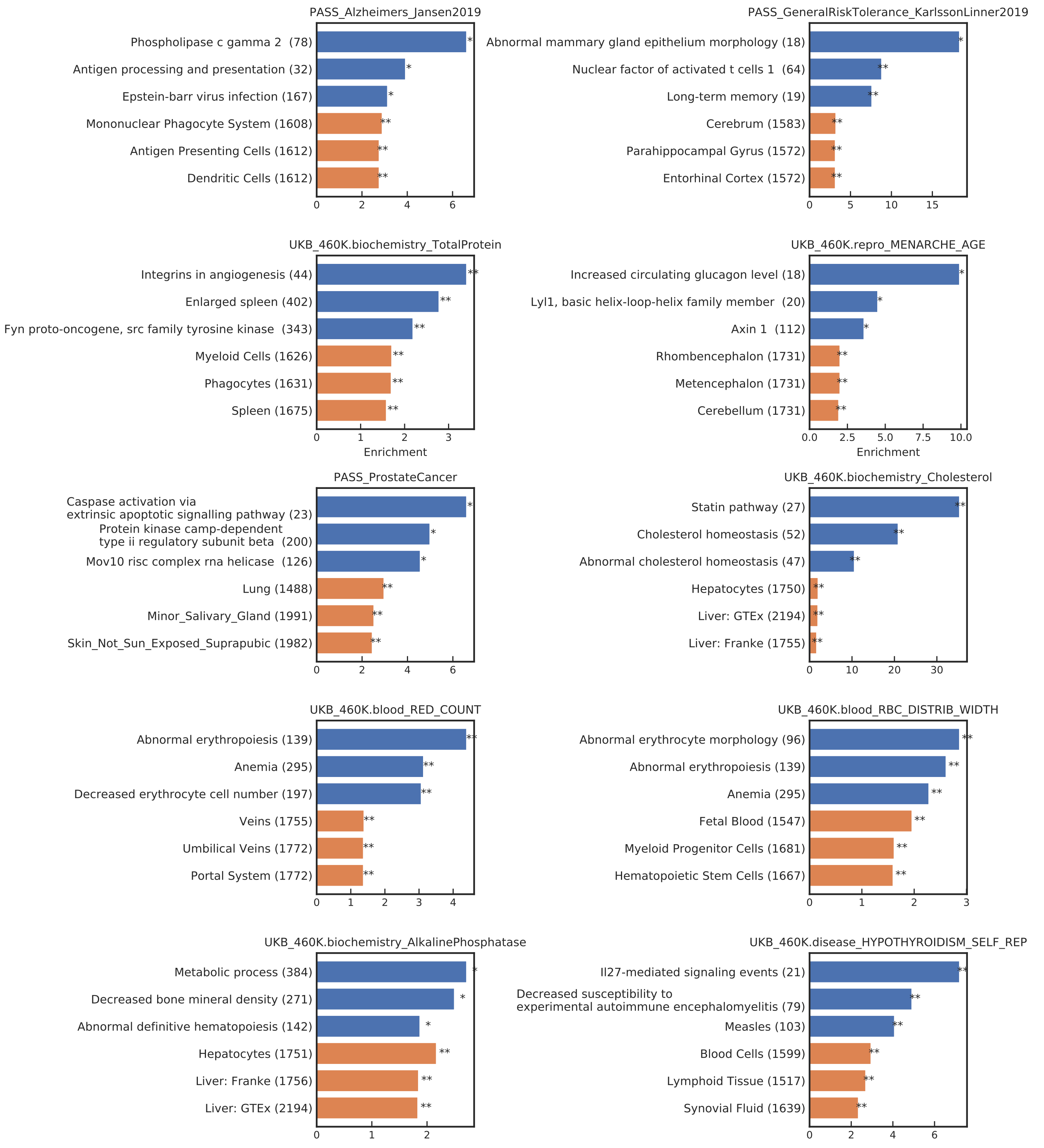

**Figure S15: Numerical GCSC results for biological pathways and specifically expressed gene sets for 10 of 43 diseases/traits.** Enrichment of gene sets in 4 traits with respect to the background of all protein coding genes. The top 3 most significant enrichment values from the specifically expressed gene analysis (in orange) and from the single trait gene set enrichment analysis (in blue) are displayed here, sorted by enrichment value. Error bars denote plus/minus one standard error. Number in parenthesis is the number of genes with a significant heritability of gene expression in at least one tissue. ** indicates passing the 20% FDR threshold and * indicates passing the 5% threshold. Gene sets pathways with the same descriptor are from different source (see supplementary table X).

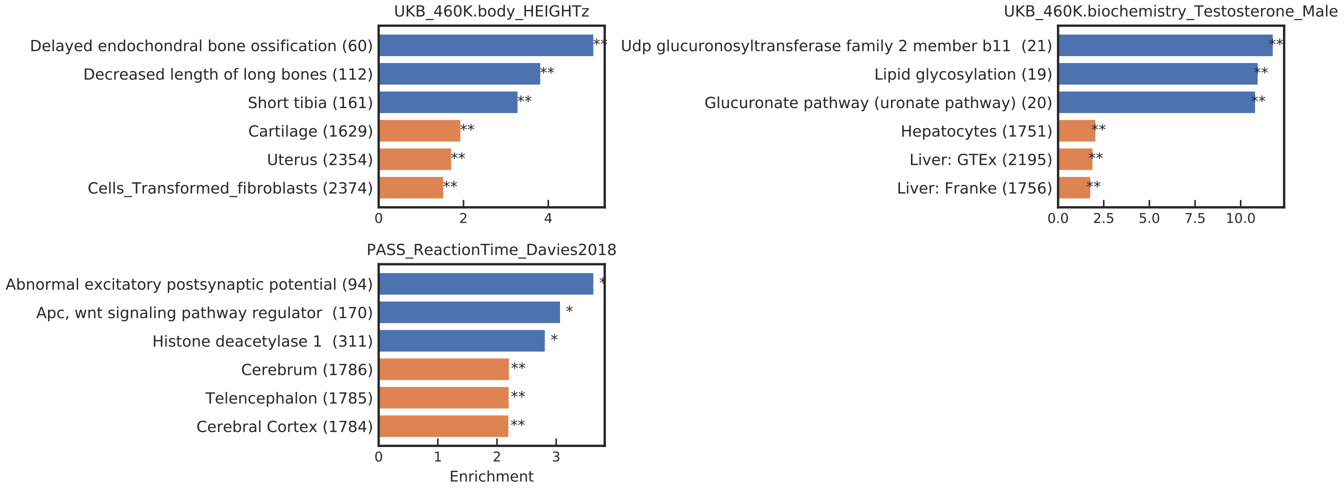

**Figure S16: Numerical GCSC results for biological pathways and specifically expressed gene sets for 3 of 43 diseases/traits.** Enrichment of gene sets in 4 traits with respect to the background of all protein coding genes. The top 3 most significant enrichment values from the specifically expressed gene analysis (in orange) and from the single trait gene set enrichment analysis (in blue) are displayed here, sorted by enrichment value. Error bars denote plus/minus one standard error. Number in parenthesis is the number of genes with a significant heritability of gene expression in at least one tissue. ** indicates passing the 20% FDR threshold and * indicates passing the 5% threshold. Gene sets pathways with the same descriptor are from different source (see supplementary table X).

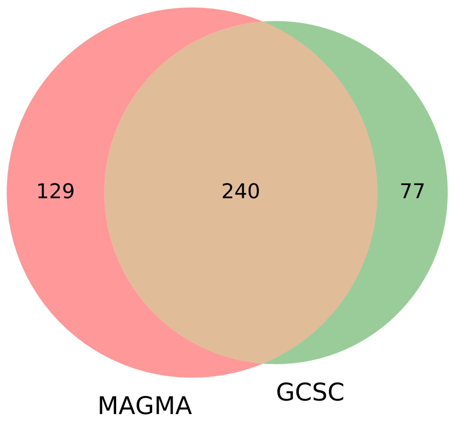

**Figure S17: Results of GCSC vs. MAGMA for non-disease-specific gene sets.** Venn Diagram of the number of gene set/trait coefficients that are positive and have a p-value less than 0.05 for MAGMA and for GCSC. A one-sided test for enrichment was used for GCSC to properly compare to MAGMA's one-sided test.

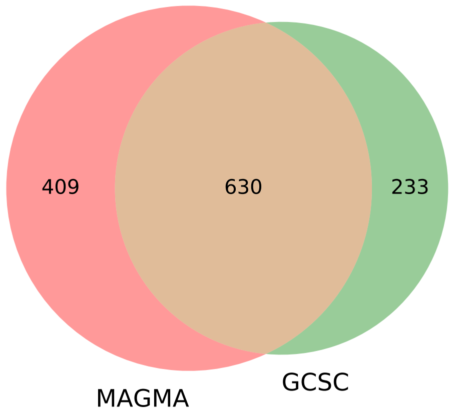

**Figure S18: Results of GCSC vs. MAGMA for specifically expressed gene sets.** Venn Diagram of the number of gene set/trait coefficients that are positive that passed a 20% FDR for MAGMA and for GCSC. A one-sided test for enrichment was used for GCSC to properly compare to MAGMA's one-sided test.

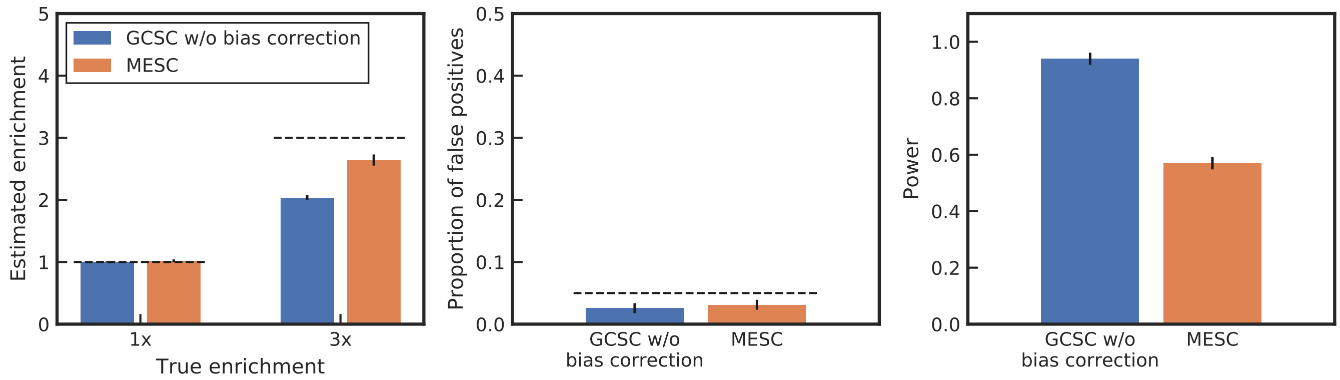

**Figure S19: Performance of GCSC with no bias correction compared to MESC.** Performance of GCSC without bias correcting the co-regulation score and MESC on simulated data. The GWAS sample size of 50,000 was used. Two tissues were simulated, one with a sample size of 80, and the other with 500. Both tissues had a fixed $h^2_{ge} = .12$ and each tissue explained 10% of trait variance. To model sharing of eQTLs between tissues, eQTL effect sizes has a correlation of 75% between tissues. In Panel (A), the estimated enrichment is plotted in simulations where there is no heritability enrichment in the gene set (1x) or when there is a 1.3x enrichment in the gene set (1.3x). In Panel (B) the proportion of 3x simulations that each method reported an enrichment p-value less than 0.05 is shown. In Panel (C), the proportion of 1x simulations in which an enrichment p-values less than 0.05 is shown. Error bars represent 1 standard error.

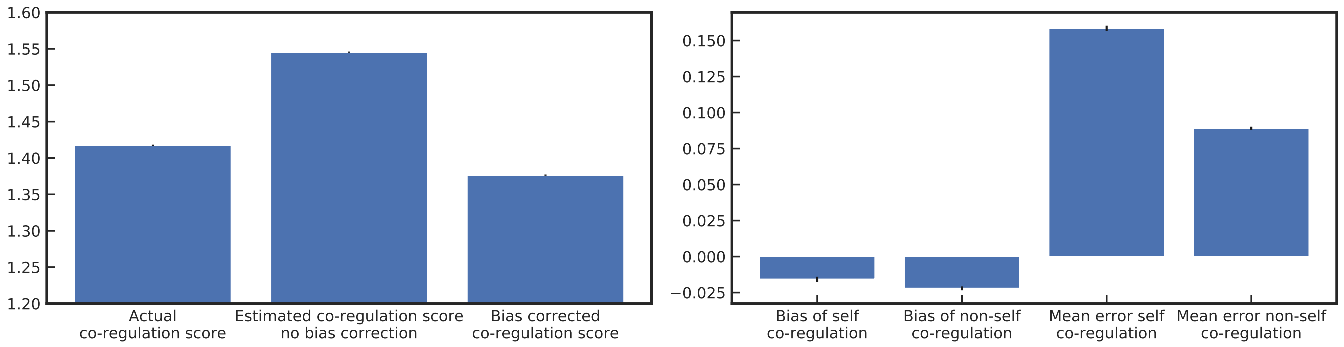

**Figure S20: Bias and error of co-regulation score estimates.** The left panel shows the mean value of the true co-regulation score, the estimated co-regulation score without bias correction and the GCSC bias corrected co-regulation score in simulations. The right panel shows the bias and error of the estimate of a gene's co-regulation score with itself only (after bias correction) and a gene's co-regulation score only with other genes, in simulations. Error is defined as the absolute value of the difference between the the estimated (and bias-corrected) value and the true value. Bias is the difference between the two values. In this figure, actual co-regulation is defined as the correlation between a gene's predicted expression and the other gene's actual gene expression to match the definition of the estimand. Error bars represent 1 standard error on the mean.

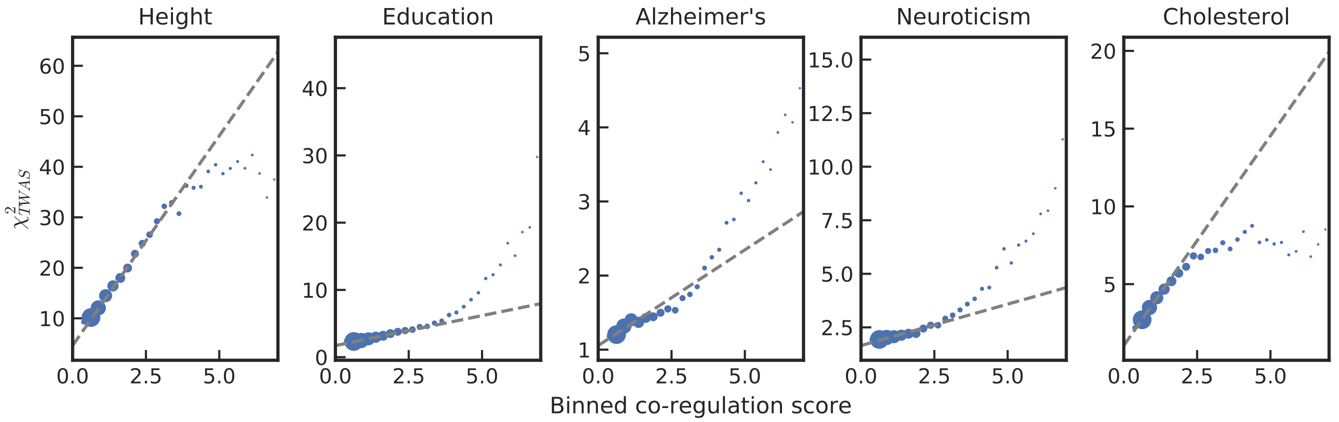

**Figure S21: Visualization of co-regulation vs TWAS $\chi^2$ for real traits.** TWAS $\chi^2$ plotted against all genes co-regulation scores for 5 traits. Grey dotted line shows result of GCSC regression. Genes were binned by co-regulation scores, an the point size indicates the sum of the regression weights of all genes in that bin.

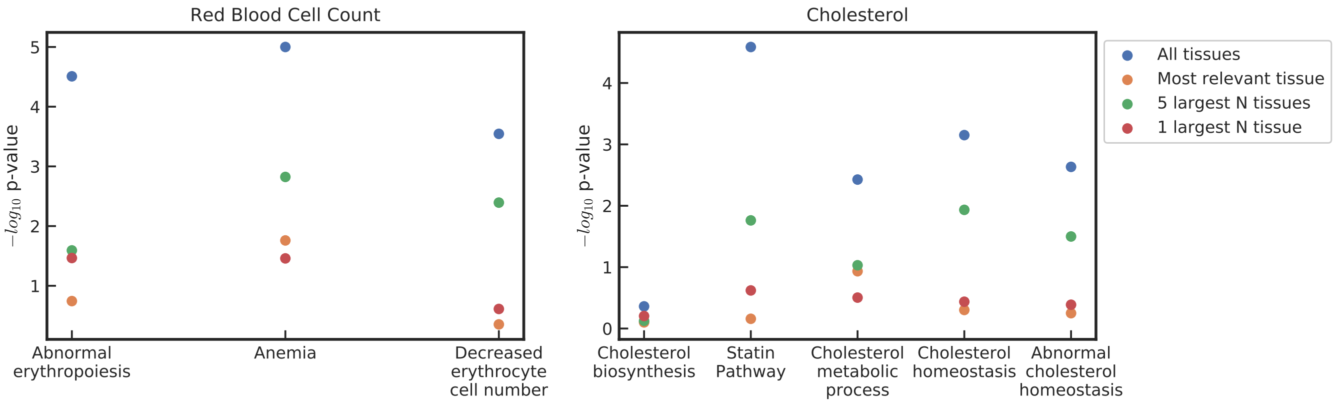

**Figure S22: Comparison of GCSC results on true positive gene sets using different tissues.** GCSC results on putative true positive gene sets for two traits. "All tissue" is GCSC using all GTExv7 tissues, "Top 5 N Tissues" uses the 5 tissues with the largest GTExv7 sample size, "Top 1 Tissue" uses just Muscle Skeletal and "Most relevant tissue" uses Liver for Cholesterol and Whole Blood for Red Blood Cell count.

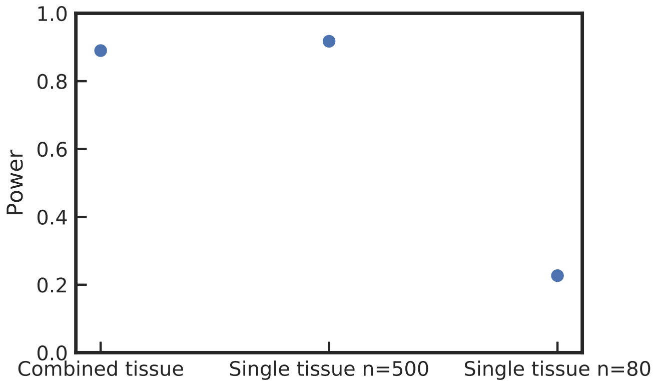

**Figure S23: Comparison of single versus multiple tissue GCSC results in simulations.** Power of GCSC in simulations with default parameters using both tissues (multi-tissue GCSC) or single-tissue GCSC using only the higher sample size tissue (n=500) or only the lower sample size tissue (n=80).

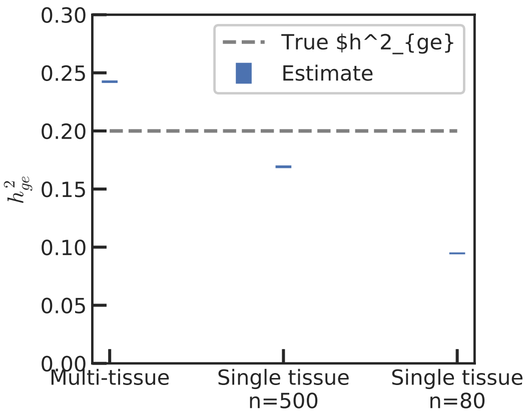

**Figure S24: Bias of $h^2_{ge}$ estimates.** Mean estimate of $h^2_{ge}$ in simulations with default parameters and two causal tissues that each mediates 10% of heritability. We ran GCSC with either both tissues or single-tissue GCSC. Single tissue GCSC $h^2_{ge}$ estimates can be above the heritability mediated by that single tissue because heritability mediated by the regression tissue can tag mediated heritability in the other tissue due to correlation in gene expression between tissues.

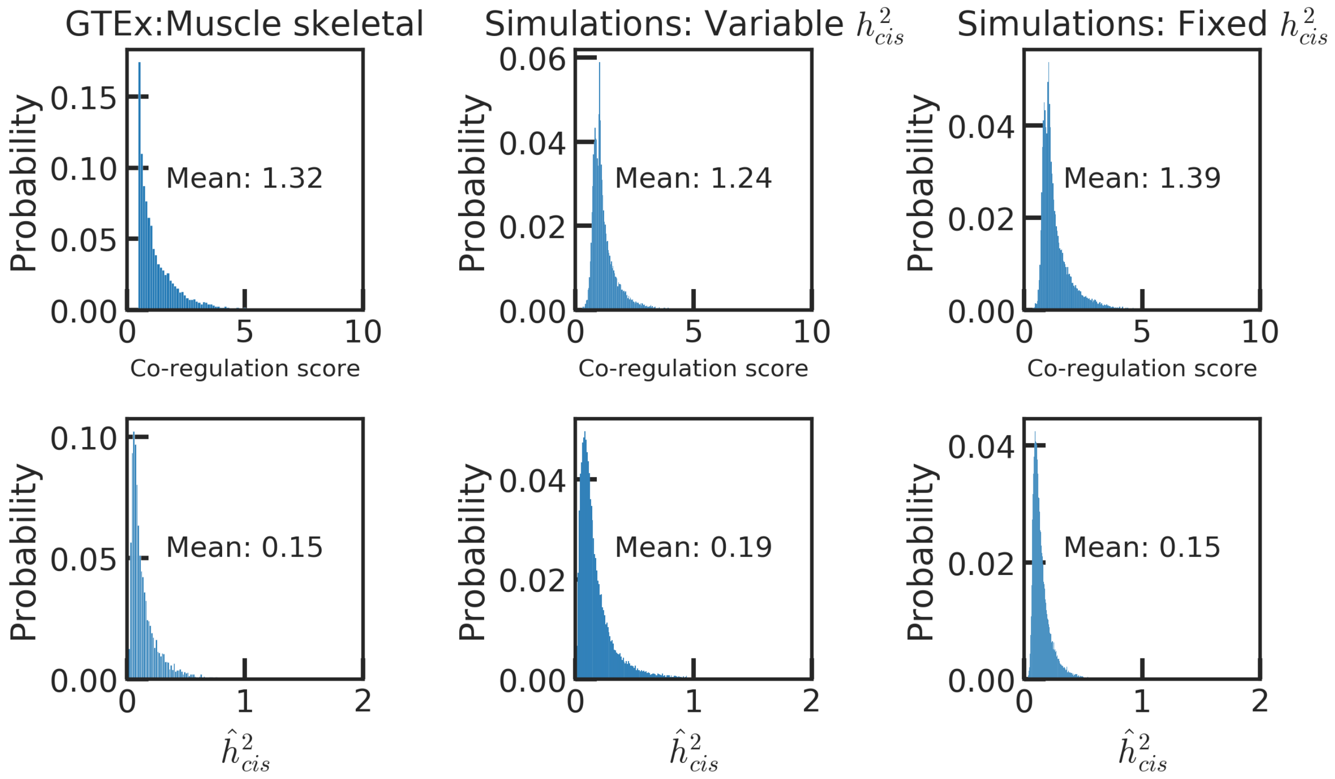

**Figure S25:** [$h^2_{cis}$ **and co-regulation score distributions in simulations and real data.** Probability density of co-regulation scores (top row) and the GCTA-estimated cis-heritability of gene expression(bottom row) in real data (left column), simulations where $h^2_{cis}$ is drawn from a distribution(middle column), and simulations where $h^2_{cis}$ is constant. The expression panel sample size was 500 in the simulations, rougly matching the Muscle skeletal sample size of 491 individuals. Note that the GCTA-estimated $h^2_{cis}$ does not usually match the real value of $h^2_{cis}$ because of estimation error.

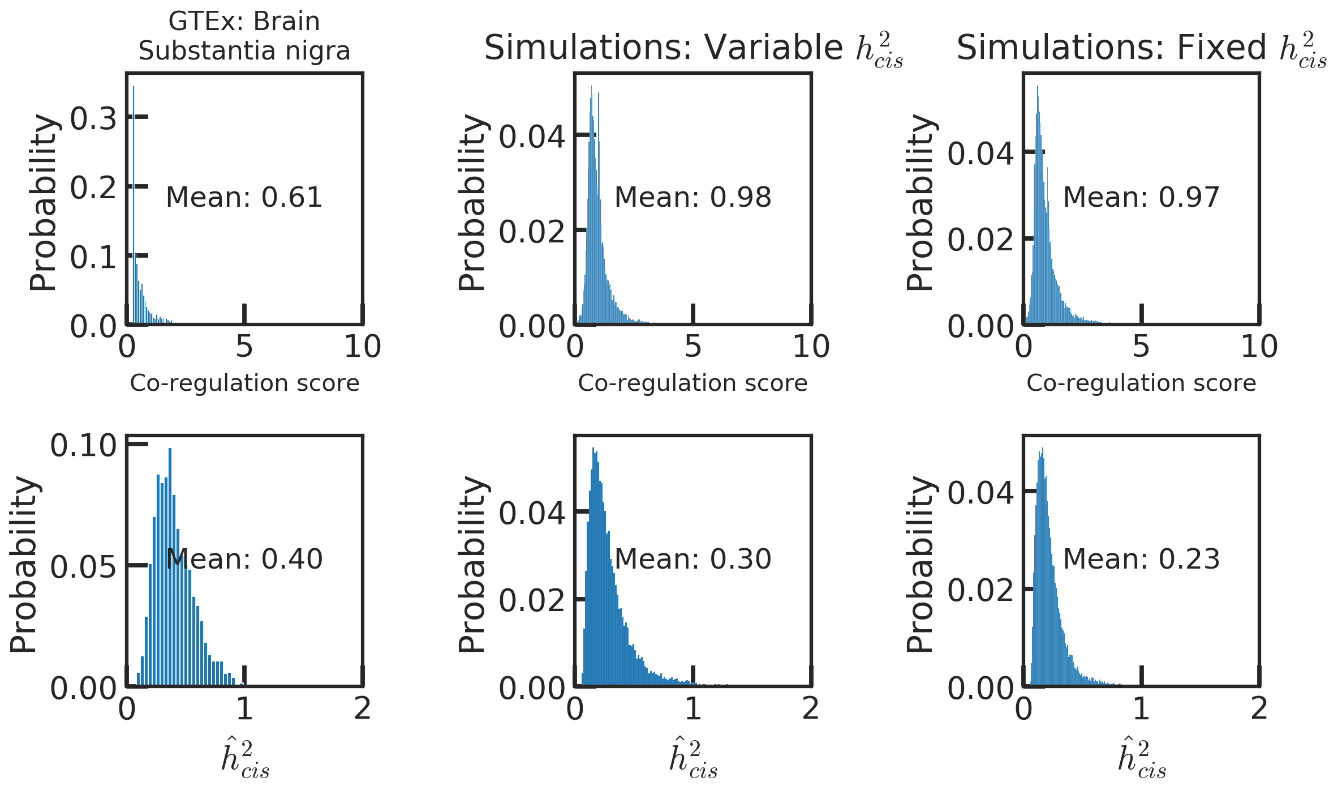

**Figure S26: $h^2_{cis}$ and co-regulation score distributions in simulations and real data.** Probability density of co-regulation scores (top row) and the GCTA-estimated cis-heritability of gene expression(bottom row) in real data (left column), simulations where $h^2_{cis}$ is drawn from a distribution(middle column), and simulations where $h^2_{cis}$ is constant. The expression panel sample size was 80 in the simulations, matching the Brain substantia nigra sample size Note that the GCTA-estimated $h^2_{cis}$ does not usually match the real value of $h^2_{cis}$ because of estimation error.

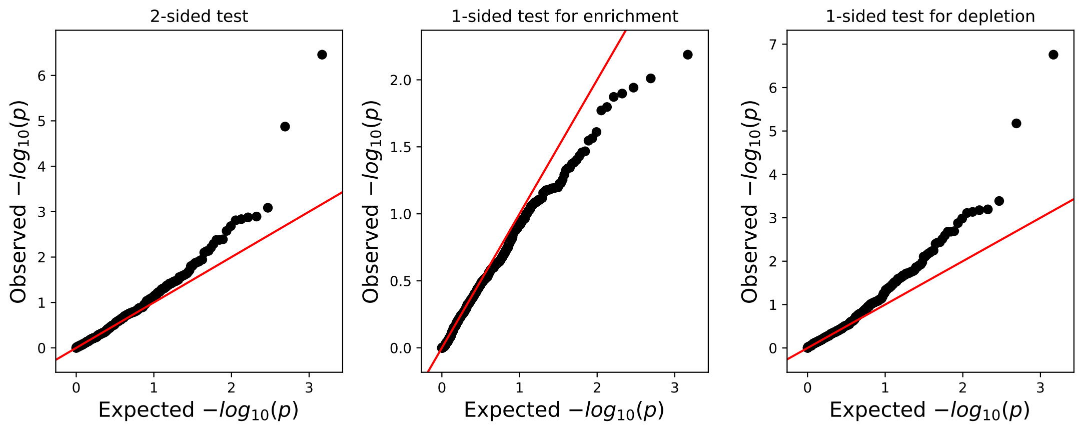

**Figure S27: QQ plot of GCSC p-values in null simulations.** QQ plots of GCSC p-values in null simulations with default parameters.

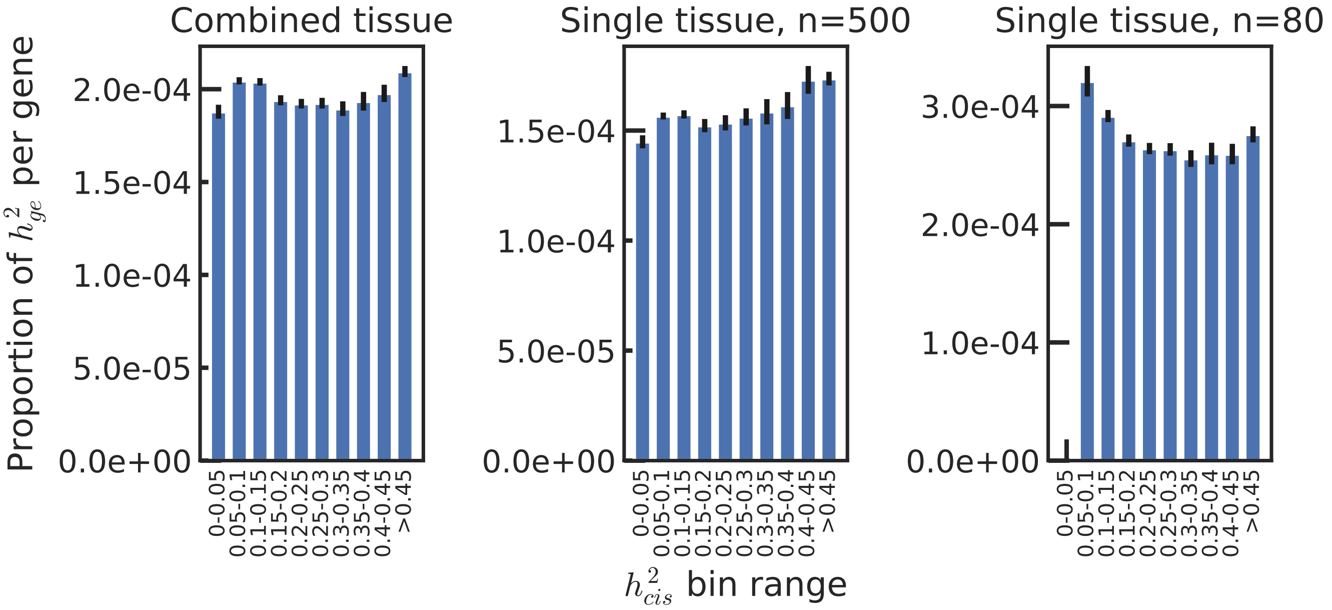

**Figure S28: $h^2_{cis}$ dependent architecture in simulations with $h^2_{cis}$ independent of $h^2_{ge}$.** Per-gene heritability explained by predicted gene expression in simulations, by $h^2_{cis}$, when $h^2_{cis}$ is independent of $h^2_{ge}$. Two tissues of sample sizes 500 and 80 were simulated and GCSC was performed using both tissues, or just one. Error bars denote +/- one standard error. Both $h^2_{cis}$ and gene to trait effect sizes were drawn from a distribution (see Methods).

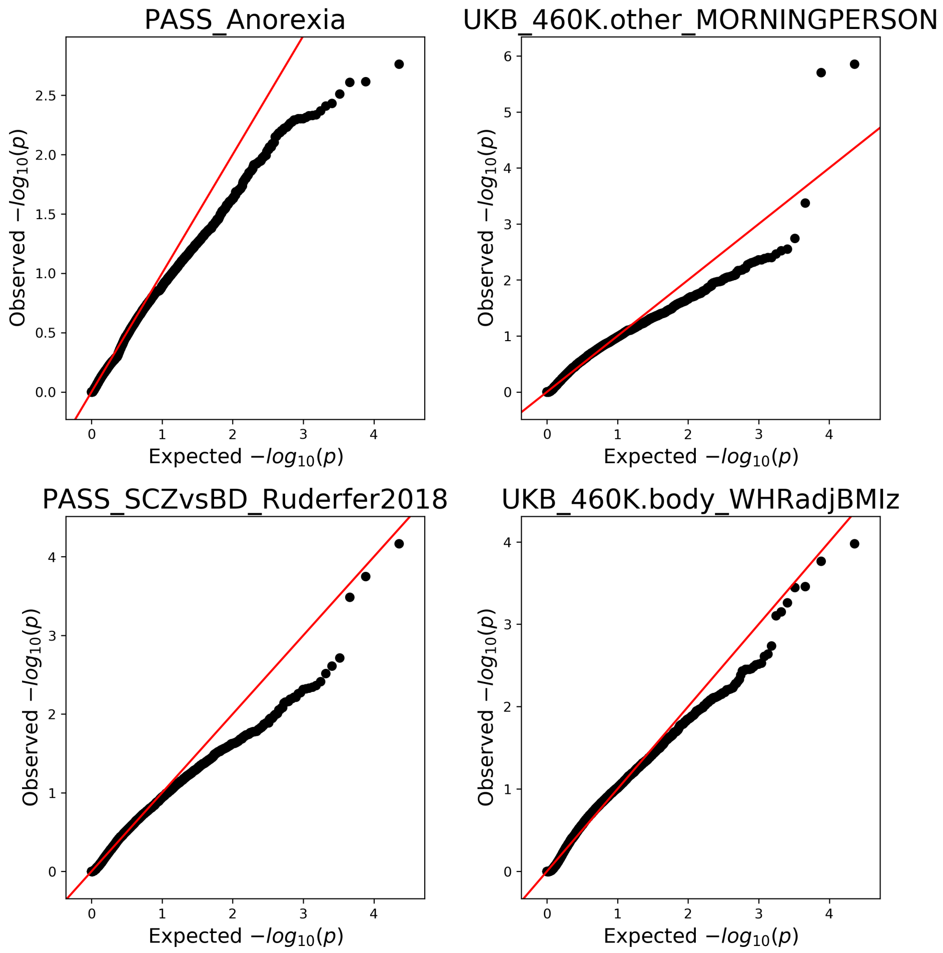

**Figure S29: QQ plot on one-sided p-values in real data.** QQ plot showing one-sided p-values for positive enrichment when applied to 11,398 gene sets from Kim et al.

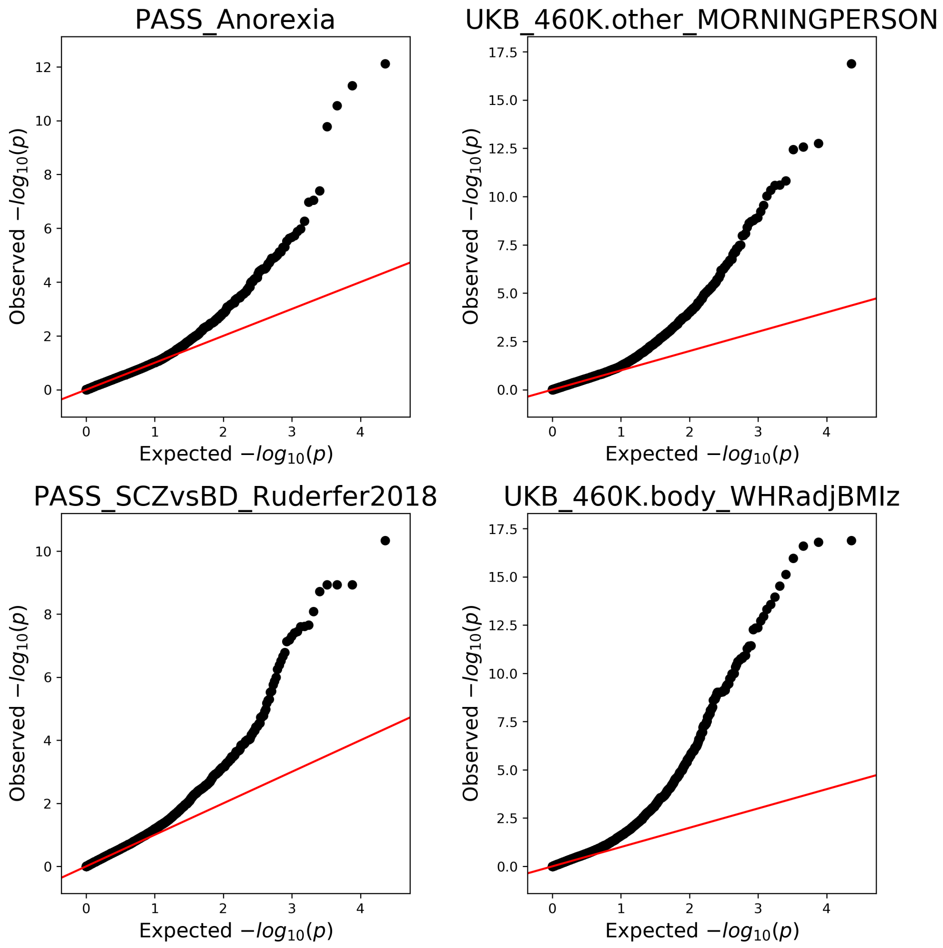

**Figure S30: QQ plot on two-sided p-values in real data.** QQ plot showing two-sided p-values when applied to 11,398 gene sets from Kim et al. A two sided p-value was used.
